# Insulin/IGF Signaling Regulates Presynaptic Glutamate Release in Aversive Olfactory Learning

**DOI:** 10.1101/2022.02.14.480437

**Authors:** Du Cheng, James Lee, Maximillian Brown, Margaret S. Ebert, Masahiro Tomioka, Yuichi Iino, Cornelia I Bargmann

## Abstract

Information flow through neural circuits is continuously modified by context-dependent learning. In the nematode *Caenorhabditis elegans,* pairing specific odors with food deprivation results in aversion to the odor. Here we identify cell-specific mechanisms of insulin/IGF receptor signaling that integrate sensory information with food context during aversive olfactory learning. Using a conditional allele of the insulin/IGF receptor DAF-2, we show that aversive learning to butanone, an odor sensed only by the AWC^ON^ olfactory neuron, requires DAF-2 in AWC^ON^. Learning requires an axonally-localized DAF-2c isoform and the insulin receptor substrate (IRS) protein IST-1, but is partly independent of the FoxO transcription factor DAF-16. Upon food deprivation, the unconditioned stimulus for learning, DAF-2 expression increases post-transcriptionally through an insulin- and *ist-1-*dependent process. Aversive learning suppresses odor-regulated glutamate release from AWC^ON^ in wild-type animals but not in *ist-1* mutants, suggesting that localized insulin signaling drives presynaptic depression to generate an aversive memory.

## INTRODUCTION

Insulin and Insulin-like Growth Factors (IGFs) are central regulators of glucose uptake, cell and organ growth, cell proliferation, and reproduction (Tokarz, et al., 2018; Yakar & Adamo, 2012). Mammalian insulin/IGF receptors are transmembrane tyrosine kinases that initiate signaling after ligand binding by cross-phosphorylation and docking of Insulin Receptor Substrate (IRS) proteins (Kavran, et al., 2014). Phosphorylated IRS proteins then recruit PI3K (Phosphatidylinositol-4,5-bisphosphate 3-kinase) to the cell membrane, where it converts the lipid PIP2 (Phosphatidylinositol 4,5-bisphosphate) to PIP3 (Phosphatidylinositol (3,4,5)- trisphosphate). PIP3 in turn activates PDK and AKT kinases, resulting in phosphorylation of multiple downstream targets. For IGF receptor signaling, important targets include FoxO, a transcription factor that is phosphorylated and retained in the cytoplasm in an inactive form by Insulin/IGF signaling. For insulin receptor signaling, critical downstream targets include a cytoplasmic signaling pathway that regulates trafficking of the plasma membrane glucose transporter GLUT-4 to support glucose uptake.

Insulin/IGF signaling also functions in learning and memory. In the nematode worm *Caenorhabditis elegans,* it regulates aversive learning and memory in contexts where food deprivation acts as an unconditioned stimulus that drives learning (Cho, et al., 2016; Tomioka, et al., 2016; Lin, et al., 2010; Tomioka, et al., 2006). In *Drosophila,* insulin/IGF signaling acts at multiple times and places to promote olfactory associative learning: it supports development of the mushroom bodies, increases learning efficiency, and drives glucose mobilization for long-term memory formation (de Tredern, et al., 2021; Chambers, et al., 2015; Naganos, et al., 2012). Mammalian insulin/IGF signaling has been implicated in CNS plasticity and learning in the hypothalamus, hippocampus, olfactory bulb, nucleus accumbens, ventral tegmental area, and infralimbic cortex (Reviewed in (Ferrario & Reagan, 2018; Dyera, et al., 2016)). Despite the robust circuit-level effects of insulin/IGF in mammals, the relevant cellular mechanisms of insulin/IGF action are only partly understood.

In *C. elegans,* pairing odors, salts, or temperature with differing food contexts can result in behavioral changes that are highly specific to the conditioned stimulus. For example, pairing the attractive odor butanone with food deprivation decreases attraction to butanone, without suppressing chemotaxis to other attractive odors (Colbert & Bargmann, 1995), whereas pairing butanone with food increases attraction to butanone (Torayama, et al., 2007). In each case, butanone attraction behavior relies upon a single olfactory neuron, AWC^ON^ (Torayama, et al., 2007; Wes & Bargmann, 2001). Similarly, pairing salt with food deprivation suppresses salt attraction, and pairing salt with food increases salt attraction (Kunitomo, et al., 2013), in both cases through a single sensory neuron, ASER. The insulin-like neuropeptide INS-1 is a candidate to convey the unconditioned stimulus of food deprivation to these sensory neurons during learning. Aversive learning after pairing food deprivation with odors or salt is lost in *ins-1* mutants (Tomioka, et al., 2016; Lin, et al., 2010; Tomioka, et al., 2006), but their appetitive learning after pairing butanone with food is normal (Lin, et al., 2010). INS-1 is released from AIA integrating neurons, and possibly other cells, and acts on sensory neurons to support aversive learning and memory. Both INS-1 and AIA are required in adults near the time of memory formation and retrieval (Cho, et al., 2016; Lin, et al., 2010; Tomioka, et al., 2006).

In addition to learning, *C. elegans* insulin/IGF signaling regulates development, metabolism, fertility, and lifespan (Murphy & Hu, 2013). Its function has been best characterized in dauer formation, a stress-induced developmental arrest. A single gene, *daf-2,* is the ortholog of both insulin receptor and IGF receptors in mammals. Like those receptors, it is predicted to be a receptor tyrosine kinase that is regulated by insulin-like peptides, which can be either agonists or antagonists of *daf-2*. *daf-2* acts through *age-1,* which encodes a PI3K homolog, and the protein kinase homologs encoded by *pdk-1, akt-1,* and *akt-2*. The primary target of this signaling pathway is the forkhead FoxO transcription factor encoded by *daf-16,* which is sequestered in the cytoplasm by *daf-2* signaling, preventing access to its transcriptional targets. Surprisingly, the *daf-2/daf-16* pathway is only modestly affected by inactivating the sole *C. elegans* IRS homolog, *ist-1* (Wolkow, et al., 2002).

Studies of salt chemotaxis learning in ASER have revealed a second *daf-2* signaling pathway that is different from the *daf-2/daf-16* transcriptional pathway (Tomioka, et al., 2016; Ohno, et al., 2014). *daf-2* encodes at least seven alternative protein isoforms, with DAF-2a being the most widely expressed and best characterized. In ASER, however, aversive learning requires a DAF-2c isoform that includes 82 additional amino acids encoded by an alternative exon (Ohno, et al., 2014). While the canonical DAF-2a protein is limited to the ASER cell body, the DAF-2c isoform required for learning is also present in the ASER axon, where its expression is increased by food deprivation (Ohno, et al., 2014). This localization suggests that axonal DAF-2c signaling may contribute to learning, and indeed, acute local activation of PI3K in the ASER axon can suppress salt attraction.

The insulin/IGF receptor *daf-2* is broadly expressed in neurons and non-neuronal tissues (Kimura, et al., 2011; Hunt-Newbury, et al., 2007). Null mutants in *daf-2* are developmentally arrested or sterile, with widespread metabolic changes, creating challenges for studying the insulin receptor pathway at cellular resolution (Murphy & Hu, 2013). Here we demonstrate that *C. elegans ist-1* mutants lacking Insulin Receptor Substrate (IRS) activity are defective in aversive olfactory learning. By characterizing *ist-1*, which is expressed only in a small number of neurons, along with single-cell knockouts of the endogenous *daf-2* locus, we identify an insulin/IGF signaling pathway required for aversive learning and show that it is acutely modulated by food deprivation, *ins-1,* and *ist-1*. The insulin/IGF receptor pathway suppresses odor-regulated glutamate release at olfactory synapses, providing a cellular mechanism for aversive memory.

## RESULTS

### The insulin receptor substrate mutant *ist-1* is defective in aversive learning

During a targeted mutant screen, we made the serendipitous finding that an *ist-1* mutation caused profound defects in aversive olfactory learning (see Methods). As learning and memory can engage different molecular mechanisms depending on training and testing protocols, we characterized behavioral responses across a range of conditions designed to distinguish between aversive learning, appetitive learning, and non-associative desensitization. In aversive learning, an Unconditional Stimulus (US), food deprivation, is paired with a Conditional Stimulus (CS), butanone exposure, resulting in reduced chemotaxis to butanone (tested at a 1:1000 dilution) (Figure 1A-B). In appetitive learning, the US is bacterial food, paired with the same CS of butanone exposure; this training results in enhanced attraction to butanone (tested at a 1:10 dilution) (Arey, et al., 2018; Torayama, et al., 2007). Finally, after exposure to very high butanone concentrations, the AWC^ON^ neuron becomes desensitized, regardless of food context, also resulting in reduced chemotaxis to 1:1000 butanone (Figure 1A-B) (Cho, et al., 2016; Pereira & Van der Kooy, 2012). By varying the amount of butanone used in training, we found that desensitization required 100-fold higher butanone training concentrations than aversive learning, and >1000-fold higher butanone training concentrations than appetitive learning (Figure 1A-B).

**Figure 1.**
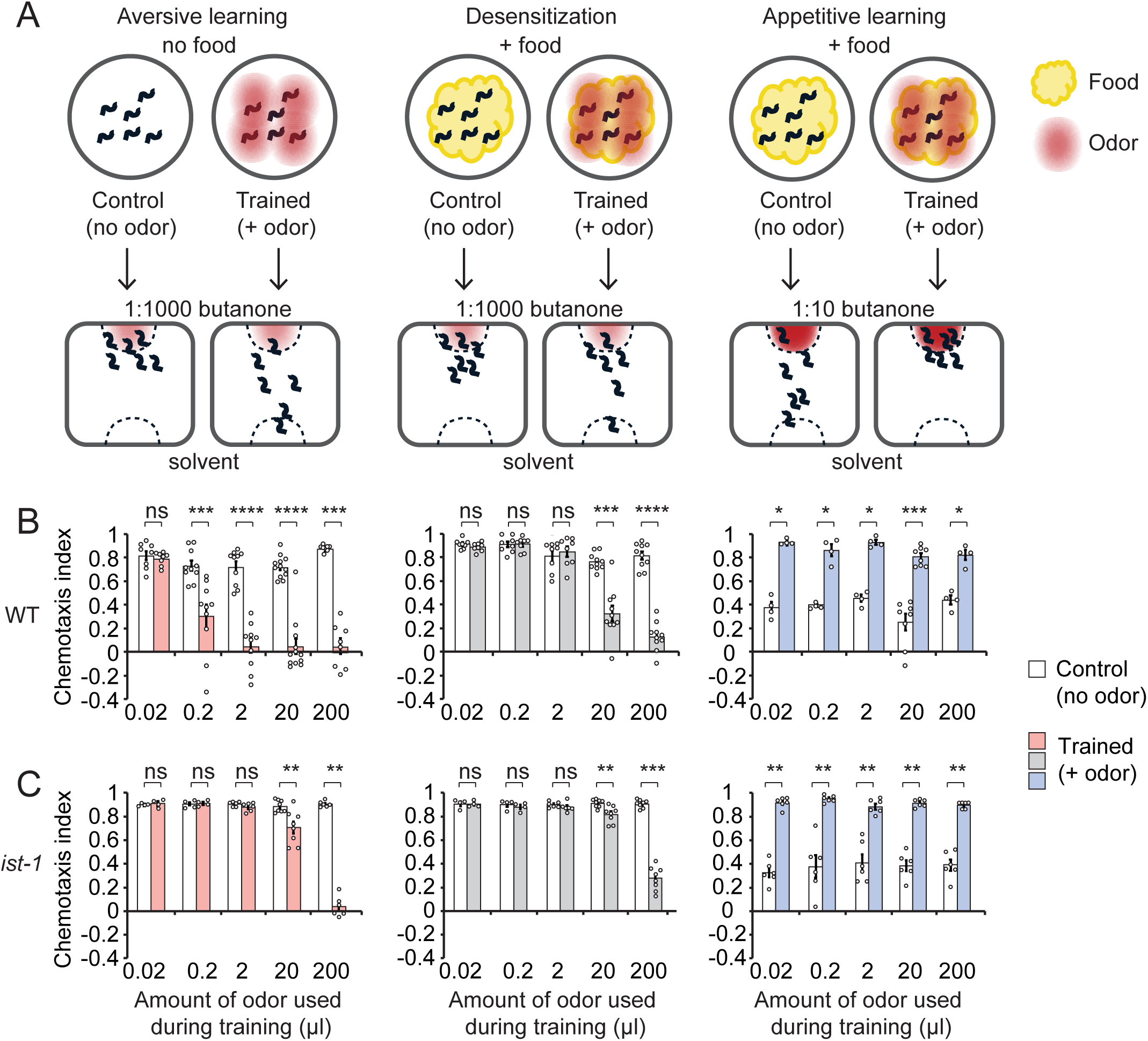
*ist-1* mutants are defective in aversive learning. A) Training and testing conditions distinguish three forms of odor learning. Associative learning to butanone can be aversive, resulting in reduced chemotaxis, or appetitive, resulting in enhanced chemotaxis, depending on whether food is present during training. Non-associative desensitization occurs after exposure to high butanone concentration regardless of the presence of food. B,C) Chemotaxis index of wild-type (B) and *ist-1* mutant (C) animals after training as in panel (A). The chemotaxis index is defined as (# of animals at odor)-(# of animals at ethanol control)/ (total # of animals in assay) after one hour. White bars, Control: animals were either deprived of food or fed for 90 min, without exposing to odor, then tested for chemotaxis to 1:1000 butanone or 1:10 butanone. Pink bars, Aversive learning: animals were deprived of food while exposed to varying concentrations of butanone odor for 90 min, then tested for chemotaxis to 1:1000 butanone. Grey bars, Desensitization: animals were fed while exposed to varying concentrations of butanone odor for 90 min, then tested for chemotaxis to 1:1000 butanone. Blue bars, Appetitive learning: animals were fed while exposed to varying concentrations of butanone odor for 90 min, then tested for chemotaxis to 1:10 butanone. Error bars indicate S.E.M. **, values differ at p ≤ 0.01. ***, values differ at p ≤ 0.001. ****, values differ at p ≤ 0.0001. ns, values are not significantly different. Statistical tests are described in Table S2.

Using these assays, we found that *ist-1* null mutants had normal chemotaxis to butanone at baseline, but were highly defective in aversive learning across a 100-fold range of butanone training conditions (Figure 1C). They had a partial defect in desensitization and were normal in appetitive learning. We further examined their behavior by using computational video tracking to examine wild-type and mutant chemotaxis in butanone gradients. *C. elegans* approaches attractants using a biased random walk, in which it makes sustained forward runs when odor increases, but reorients its movement when odor decreases (Figure S1A) (Levy & Bargmann, 2020; Pierce-Shimomura, et al., 1999). These reorientation behaviors were similar in naïve wild-type and *ist-1* animals (Figure S1B). After training, reorientations were no longer correlated with the direction of the odor source in wild-type animals, but *ist-1* mutants retained a substantial odor-oriented reorientation bias consistent with chemotaxis (Figure S1B-C).

### *ist-1* is required in the AWC^ON^ neuron for aversive learning

The *ist-1* gene encodes a protein related to mammalian IRS proteins, with predicted phospholipid-binding pleckstrin homology (PH), phosphotyrosine binding (PTB), and Phosphatidyl Inositol 3-kinase (PI3K) binding domains (Wolkow, et al., 2002). Four *ist-1* mRNAs differ based on their initiation sites and 5’ exons (*ist-1b/e* versus *ist-1c/d)* and alternative splicing of two internal exons (*ist-1b/c* versus *ist-1d/e*) (Figure 2A). We observed significant defects in aversive learning in a newly-generated CRISPR-Cas9 allele predicted to affect only the *ist-1b/e* (long) isoforms, *ky1074*, and stronger aversive learning defects in three alleles that disrupted all isoforms, *ky1071, ky1075,* and *ok2706* (Figure 2B).

**Figure 2.**
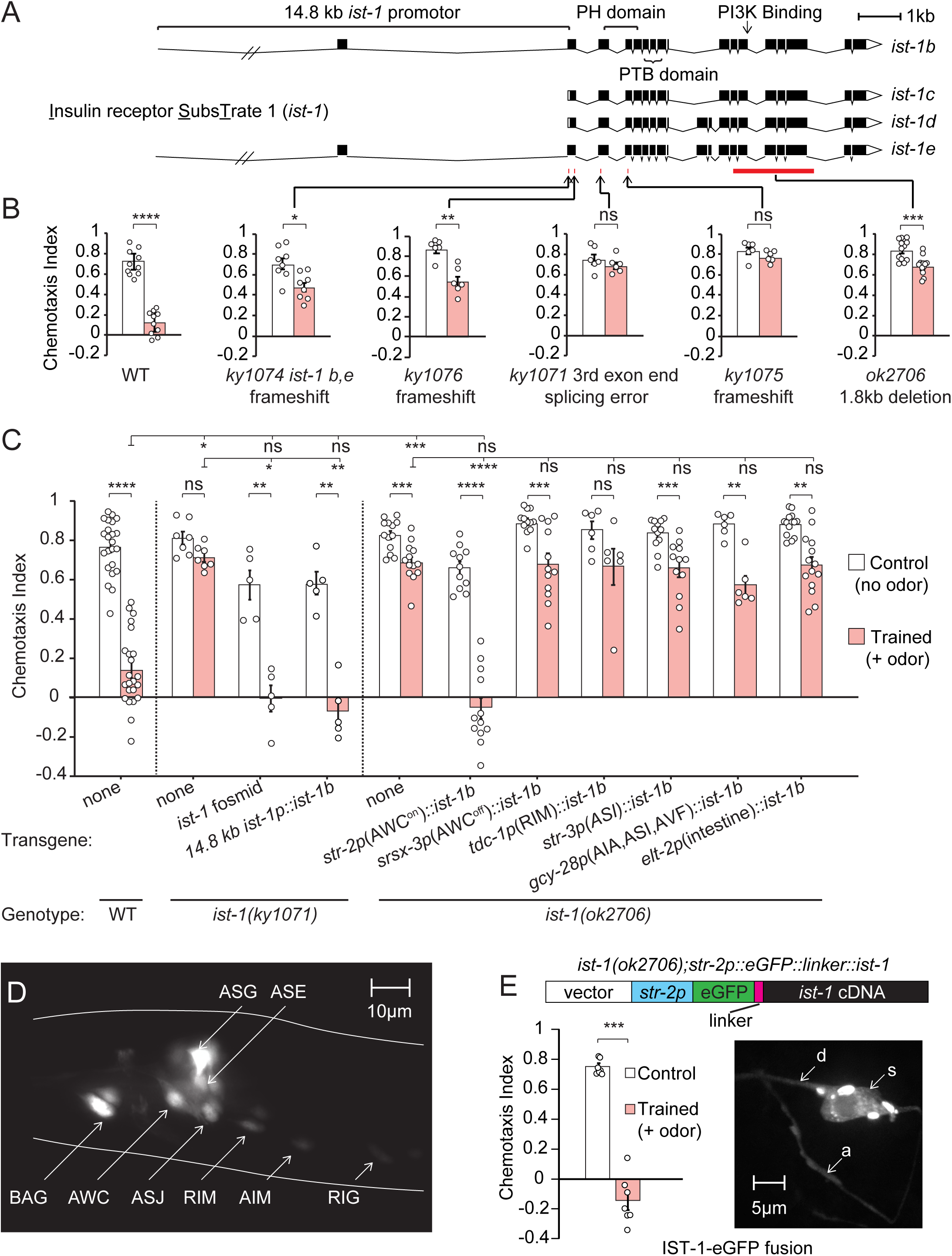
*ist-1* is required in AWC^ON^ for butanone aversive learning. A) Genomic structure of the *C. elegans* insulin receptor substrate-1 *(ist-1)* gene. Four isoforms of *ist-1* are generated from alternative promoters and alternative splicing. Solid blocks correspond to exons, and locations of mutations are indicated by a red line or a red bar. *ky1071* was a background mutation discovered in the *flp-20(pk1596)* strain. *ok2706* was obtained from the *Caenorhabditis* Genetics Center. *ky1074, ky1076* and *ky1075* were generated by targeted CRISPR/Cas-9 mutagenesis (see Methods). B) Chemotaxis and aversive learning in *ist-1* mutants. Animals were food-deprived (control, white bars) or food-deprived with 20 µl butanone odor (trained, pink bars) for 90 min and tested for chemotaxis to 1:1000 butanone. C) Transgenic rescue of *ist-1* mutants. Animals were trained and tested as above. Left, full rescue of *ky1071* mutants with a fosmid covering the entire genomic sequence of *ist-1* or a 14.8 kb promoter fragment driving an *ist-1b* cDNA. This strain included the original *flp-20* mutation. Right, rescue of *ist-1(ok2706)* mutants with the *ist-1b* cDNA under different cell-specific promoters. Only the *str-2* promoter that expresses *ist-1* in AWC^ON^ rescued aversive olfactory learning. D) Expression of an *ist-1::eGFP* reporter with 14.8 kb of genomic DNA encompassing 9.3 kb upstream of the first *ist-1* exon (the initiation site of isoforms *ist-1b* and *ist-1e)* and 5.3 kb upstream of the second exon (the initiation site of isoforms *ist-1c* and *ist-1d)*. E) Localization of an eGFP::IST-1 fusion protein expressed under the AWC^ON^ -selective *str-2* promoter. Top, diagram of the expression construct. Lower left, transgenic rescue of *ist-1(ok2706)* by *str-2::eGFP::IST-1*. Animals were trained and tested as above. Lower right, representative maximal intensity projection of of IST-1::eGFP in AWC^ON^ neuron, showing fluorescence in the dendrite (d), axon (a), and cell soma (s). Aggregated protein in the cell body is likely due to overexpression. Error bars indicate S.E.M. *, values differ at p ≤ 0.05. **, values differ at p ≤ 0.01. ***, values differ at p ≤ 0.001. ****, values differ at p ≤ 0.0001. ns, values are not significantly different. Statistical tests are described in Table S2.

Expression of an *ist-1b* cDNA under a 14.8 kb genomic fragment encompassing 9.3 kb upstream of the first initiation site, the first exon, and the 5.3 kb intron before the second initiation site resulted in full rescue of learning (Figure 2C). The same 14.8 kb fragment drove reliable GFP expression in eight pairs of head neurons (AWC, ASE, ASG, ASI, and BAG sensory neurons and RIM, AIM, and RIG interneurons (Figure 2D) and in some tail neurons (Figure S2C), overlapping the *ist-1* expression pattern obtained from single-cell RNA sequencing data (Taylor, et al., 2021). Notably, *ist-1* expression is absent in many neuronal and non-neuronal cells that express the insulin receptor *daf-2* (Kimura, et al., 2011; Hunt-Newbury, et al., 2007). To define cells in which *ist-1* acts in aversive learning, *ist-1b* was expressed under shorter regions of the endogenous *ist-1* promoter and under cell-specific promoters that overlapped with its expression pattern (Figure 2C, Figure S2). Expression of *ist-1* in the AWC^ON^ neuron, which is the primary butanone-sensing neuron, fully rescued aversive learning in *ist-1* null mutants, whereas expression in other neurons or in the intestine did not (Figure 2C, Figure S2). A GFP-tagged IST-1 protein that rescued the *ist-1* aversive learning defect was broadly distributed in the axon, dendrite and cell body of AWC^ON^ and appeared to be excluded from the nucleus (Figure 2E). These results suggest that *ist-1* functions autonomously in the AWC^ON^ sensory neuron to drive aversive learning.

### *ist-1* interacts with insulin pathway genes in aversive learning

Aversive learning to butanone requires the insulin-related gene *ins-1,* which is expressed in the AIA interneurons, other neurons, and intestine, and the *daf-2*-regulated PI3 kinase *age-1,* which is required in AWC^ON^ neurons (Takeishi, et al., 2020; Cho, et al., 2016) (Figure 3A-B). We found that several additional genes in the insulin pathway had either butanone chemotaxis defects or aversive learning defects. Loss of function mutants in the *akt-1* kinase had a deficit in aversive learning comparable to that of *ins-1, ist-1* and *age-1* mutants (Figure 3B). Insulin/IGF signaling in both vertebrates and invertebrates is negatively regulated by PTEN, a phosphatase that dephosphorylates PIP3 to form PIP2; *daf-18* PTEN phosphatase mutants failed to chemotax to butanone before or after conditioning (Figure 3A-B).

**Figure 3.**
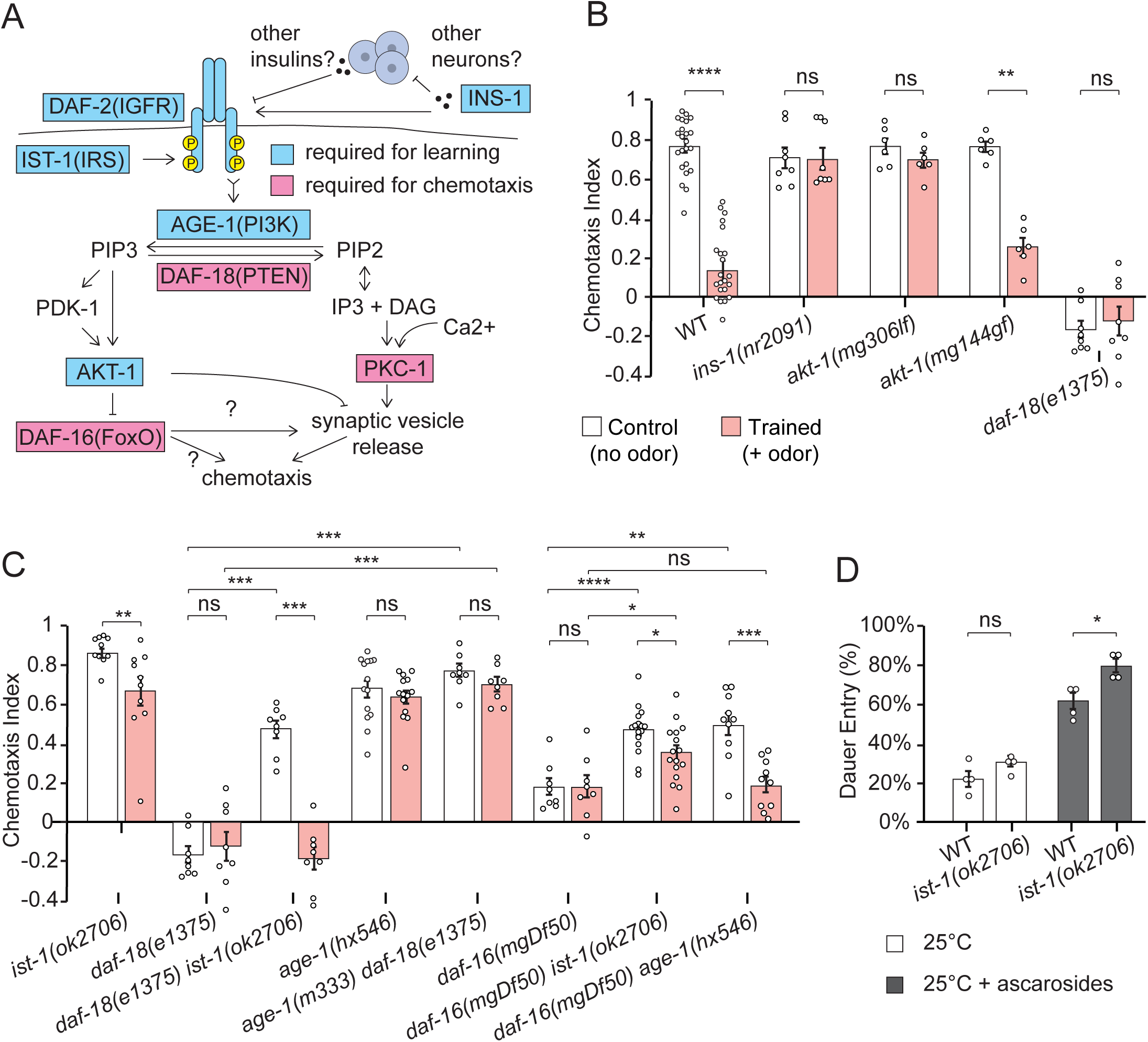
*ist-1* interacts with the insulin/IGF signaling pathway in aversive learning. A) Summary diagram of *C. elegans* insulin/IGF pathway highlighting genes that affect butanone chemotaxis (pink) or aversive learning (blue). B,C) Chemotaxis and aversive learning in insulin pathway mutants and double mutants. Animals were food-deprived (control, white bars) or food-deprived with 20 µl butanone odor (trained, pink bars) for 90 min and tested for chemotaxis to 1:1000 butanone. Defects in *ins-1, age-1,* and *daf-16* were as previously reported (Cho, et al., 2016; Daniels, et al., 2000). *daf-16(mgDf50)* and *age-1(m333)* are null alleles; *daf-18(e1375)* and *age-1(hx546)* are partial loss of function alleles. D) Dauer formation in wild-type and *ist-1(ok2706)* animals grown in mildly dauer-inducing conditions (limited food at 25°C) or strongly dauer-inducing conditions (limited food at 25°C supplemented with 73 nM each of ascarosides C3 and C6). Error bars indicate S.E.M. *, values differ at p ≤ 0.05. **, values differ at p ≤ 0.01. ***, values differ at p ≤ 0.001. ****, values differ at p ≤ 0.0001. ns, values are not significantly different. Statistical tests are described in Table S2.

To further determine the relationship between *ist-1* and other genes in the pathway, we examined double mutants. Double mutants between *ist-1* and the PTEN phosphatase *daf-18* had mixed phenotypes: the *daf-18* butanone chemotaxis defect was partially restored in *daf-18; ist-1* double mutants, and aversive learning was restored as well (Figure 3C). These results are consistent with a model based on the mammalian insulin pathway, in which the insulin receptor DAF-2 activates the PI3 kinase AGE-1, the adaptor protein IST-1 potentiates the DAF-2/AGE-1 interaction, the DAF-18 phosphatase antagonizes AGE-1, and simultaneous loss of both IST-1 and DAF-18 permits residual AGE-1 activation by DAF-2 (Figure 3A). In agreement with this model focused on AGE-1 activation, *age-1; daf-18* double mutants resembled *age-1*, with robust chemotaxis and no aversive learning (Figure 3C). However, the *daf-18* mutation used in this experiment is not a null allele, which limits the interpretation of these results.

*daf-16* FoxO mutants do not chemotax to butanone (Daniels, et al., 2000). In addition, their behavior was not modified by aversive conditioning (Figure 3C). *daf-16; ist-1* double mutants had a mixed phenotype: they had intermediate levels of chemotaxis to butanone, with diminished levels of aversive learning (Figure 3C). Since both *ist-1* and *daf-16* alleles were null alleles, this epistasis analysis indicates that each of these genes has some functions that are independent of the other. The *daf-16* null phenotype was also partly suppressed in *daf-16; age-1* double mutants (Figure 3C).

The role of *ist-1* in dauer development, the target of the canonical *C. elegans* insulin/IGF pathway, has been studied only through RNA interference (Wolkow, et al., 2002). Using the *ist-1* deletion mutant *ok2706*, we confirmed that *ist-1* mutants did not form dauers under well-fed conditions (data not shown). They were also similar to wild-type in mildly dauer-inducing conditions with limited food at high temperatures (Figure 3D), and were only slightly more likely than wild-type to form dauers when exposed to synthetic pheromones that resulted in substantial dauer formation (Figure 3D).

In summary, *ist-1* (insulin receptor substrate), *ins-1* (an insulin), *age-1* (PI3K), and *akt-1* (AKT) are all required for aversive butanone learning, and *daf-18* (PTEN) antagonizes *ist-1* in a way that is consistent with *ist-1* and *daf-18* action upstream of *age-1*. However, there are distinctions between the aversive learning pathway and the canonical *C. elegans* insulin/IGF pathway (Murphy & Hu, 2013). *ins-1* acts as an inhibitor of insulin signaling in dauer formation, but apparently as an activator (with a similar phenotype to *age-1)* in learning. *ist-1* and *akt-1* are essential for aversive learning, but not for dauer formation, where *akt-1* is redundant with *akt-2*. *daf-16* is the final output of the *daf-2* pathway in dauer formation, and is epistatic to all other genes, but *ist-1* function in aversive learning is partially independent of *daf-16*.

### The DAF-2c insulin receptor isoform supports aversive learning in AWC^ON^

One concern in studying insulin pathway mutants, particularly the essential gene *daf-2,* is that systemic physiological abnormalities in mutants might indirectly affect learning. To begin to address this point, we examined two *daf-2* mutants that do not have overt developmental defects, but have taste avoidance defects when salt is paired with food deprivation (Nagashima, et al., 2019; Ohno, et al., 2014) (Figure 4A). The *daf-2(pe1230)* missense mutation affecting the extracellular domain was highly defective for butanone aversive learning (Figure 4B). A partial aversive learning defect was caused by the DAF-2c-specific allele *daf-2c(pe2722)* (Figure 4B), which selectively disrupts isoform-specific exon 11.5 that adds 82 amino acids to DAF-2c (Nagashima, et al., 2019; Ohno, et al., 2014).

**Figure 4.**
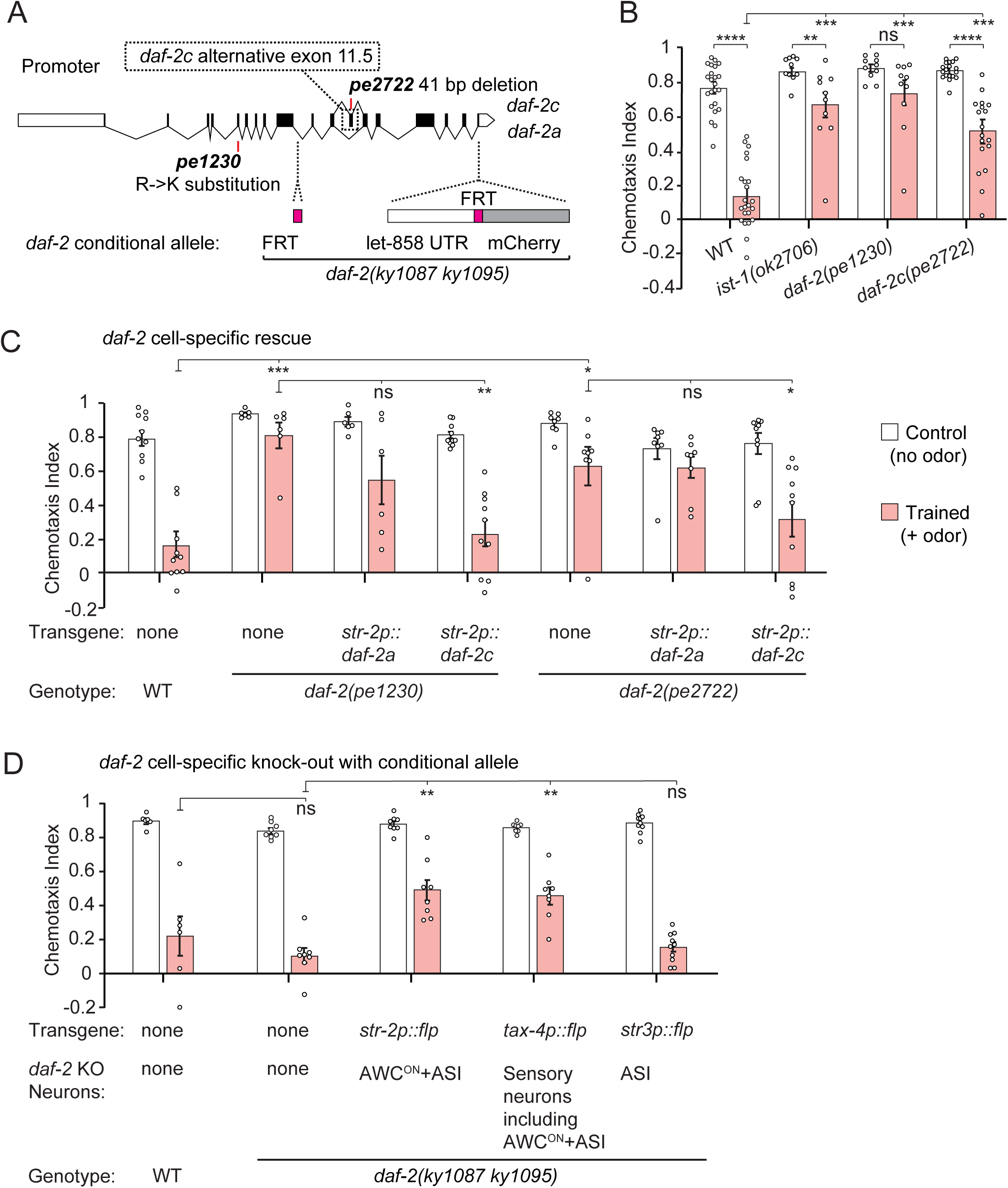
The DAF-2 insulin/IGF receptor is required in AWC^ON^ for aversive learning. A) Top, genomic structure of *daf-2* and mutations used in this study. White boxes denote promoter and 3’UTR regions; black boxes denote exons. The *daf-2a* and *daf-2c* isoforms differ by inclusion of the alternative exon 11.5. The mutations in *daf-2(pe1230)* and *daf-2c(pe2722)* both affect the extracellular domain of the predicted DAF-2 protein. To generate a conditional *daf-2* knockout allele for this study, two FRT sites flanking half of the *daf-2* gene were inserted by CRISPR-Cas9 mutagenesis of the endogenous *daf-2* gene (see Methods). B) Chemotaxis and aversive learning in wild-type, *ist-1(ok2706), daf-2(pe1230),* and *daf-2c(pe2722)* animals. Animals were food-deprived (control, white bars) or food-deprived with 20 µl butanone odor (trained, pink bars) for 90 min and tested for chemotaxis to 1:1000 butanone. C) Transgenic rescue of *daf-2(pe1230)* and *daf-2c(pe2722)* mutants with *daf-2a* or *daf-2c* cDNAs expressed under the AWC^ON^ -selective *str-2* promoter. Animals were trained and tested as above. D) Chemotaxis and aversive learning in animals with cell-selective flippase transgenes injected into the *daf-2frt* strain to generate conditional knockouts. In each strain, the site and efficiency of flippase recombination was determined using a reporter transgene (see Methods). Significant learning defects were observed upon flippase expression under the AWC^ON^-selective *str-2* promoter or the *tax-4* promoter that is expressed in AWC and nine other pairs of sensory neurons. Flippase expression under the ASI-specific *str-3* promoter did not elicit learning defects. Animals were trained and tested as above. Error bars indicate S.E.M. *, values differ at p ≤ 0.05. **, values differ at p ≤ 0.01. ***, values differ at p ≤ 0.001. ****, values differ at p ≤ 0.0001. ns, values are not significantly different. Statistical tests are described in Table S2.

Next, we tested full-length *daf-2a* and *daf-2c* cDNAs expressed under the AWC^ON^-selective *str-2* promoter for rescue of *daf-2* mutant phenotypes. *daf-2c* fully rescued the aversive learning defects of both *daf-2(pe1230)* and *daf-2c(pe2722)* mutants. *daf-2a* did not significantly rescue aversive learning in *daf-2c(pe2722)* or *daf-2(pe1230)* mutants, although there was a trend toward rescue in *daf-2(pe1230)* (Figure 4C). The mutant and cell-specific rescue results suggest that the *daf-2c* isoform has a primary role in aversive learning in AWC^ON^, although other isoforms such as *daf-2a* may also contribute.

To further refine the site of *daf-2* action, we generated a conditional allele of the endogenous *daf-2* locus susceptible to flippase-FRT inactivation. Two FRT sites flanking the second half of the *daf-2* gene were inserted by CRISPR-Cas9 recombination, and flippase was expressed under cell-selective promoters to inactivate *daf-2* only in selected cells (Figure 4A). Expression of flippase under an AWC^ON^-selective promoter resulted in a partial aversive learning defect after butanone conditioning (Figure 4D). As a flippase activity reporter indicated that this transgene was also active in ASI neurons, and occasionally in other cells (Muñoz-Jiménez, et al., 2017) (Figure 4D and Methods), we used other promoters to confirm AWC^ON^ as a relevant site of *daf-2* action. An ASI-specific *daf-2* knockout did not affect aversive learning, while a sensory-specific *daf-2* knockout that affected AWC^ON^ and other sensory neurons disrupted aversive learning. Together, these results support a role for *daf-2* in AWC^ON^ in aversive learning. The incomplete effect of *daf-2* inactivation in AWC^ON^ in may be due to *daf-2* activity in other cells, incomplete flippase recombination (see Methods), or pre-existing *daf-2* mRNA or protein that persists in AWC^ON^ after recombination.

### IST-1 has a minor role in the DAF-2c pathway for taste avoidance learning

The involvement of the *daf-2c* isoform in aversive olfactory learning suggested an analogy with taste avoidance learning, the associative learning behavior in which *daf-2c* was first characterized (Ohno, et al., 2014). Taste avoidance learning has similar properties to olfactory avoidance learning: when high salt is paired with food deprivation, animals lose their attraction to high salt (Figure 5A). *ist-1* mutants had a partial decrease in taste avoidance learning (Figure 5B), while retaining normal preferences when high or low salt was paired with food (Figure S3). This defect was rescued by expressing *ist-1* in ASER, the sensory neuron required for taste avoidance learning (Figure 5B). The *ist-1* mutant defect in taste avoidance learning was milder than the defect in *daf-2c(pe2722)* mutants. *daf-2c; ist-1* double mutants had strong defects resembling those of *daf-2c* (Figure 5B).

**Figure 5.**
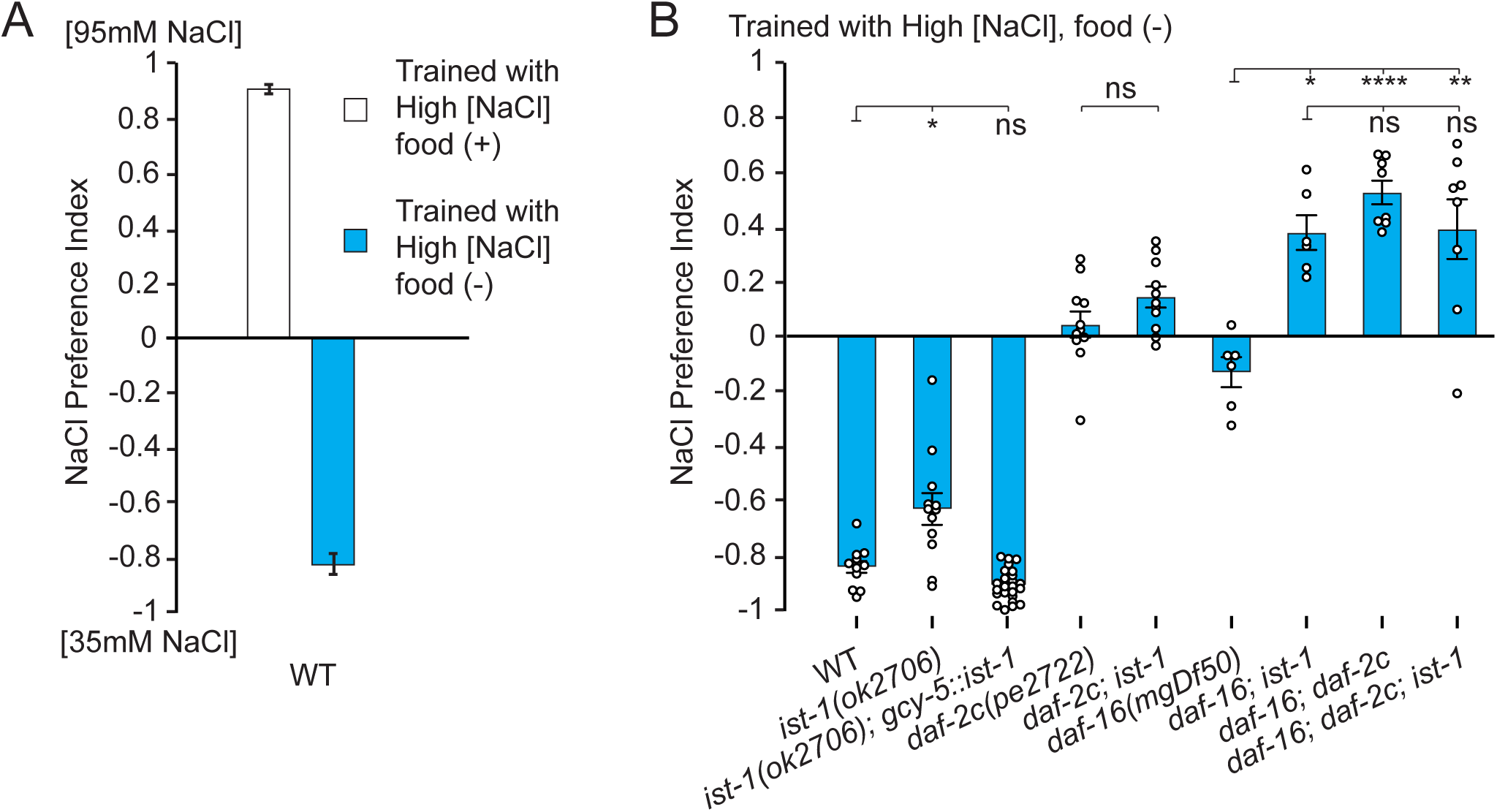
*ist-1* in ASER interacts with the insulin/IGF pathway in taste avoidance learning. A) Taste avoidance learning. Pairing food with 100 mM NaCl results in chemotaxis to high salt (95 mM) in a salt gradient (trained, white bars). Pairing food deprivation with 100 mM NaCl results in taste avoidance learning, expressed as chemotaxis to low salt (35 mM) in a salt gradient (trained, blue bars). The preference index is defined as (# of animals at high salt)-(# of animals at low salt)/ (total # of animals in assay) after one hour. B) Taste avoidance learning in insulin pathway mutants including *ist-1*, *daf-2c*, and *daf-16,* and rescue of taste avoidance learning by *ist-1* expression under the ASER-specific *gcy-5* promoter. Animals were food-deprived with 100 mM NaCl for 5 hours (trained, blue bars) and tested for chemotaxis in a salt gradient as above. Error bars indicate S.E.M. *, values differ at p ≤ 0.05. **, values differ at p ≤ 0.01. ****, values differ at p ≤ 0.0001. ns, values are not significantly different. Statistical tests are described in Table S2. Additional salt training and chemotaxis results are in Figure S3.

*daf-16* mutants are also defective in taste avoidance learning, resembling *daf-2c(pe2722)* (Figure 5B). Double mutants between the *daf-16* and *ist-1* mutants had a stronger defect than either single mutant (Figure 5B). A similar enhanced defect was observed in *daf-16; daf-2c* double mutants (Nagashima, et al., 2019) and in *daf-16; daf-2c; ist-1* triple mutants (Figure 5B). These results are consistent with overlapping functions for *daf-2c* and *ist-1*, and distinct functions for *daf-16,* in taste avoidance learning.

In summary, *ist-1, daf-2c,* and *daf-16* all affect both aversive olfactory learning and taste avoidance learning. For salt chemotaxis learning, the *daf-2c(pe2722)* mutant defect was more severe than that of *ist-1,* whereas for aversive olfactory learning the *ist-1* defect was more severe than that of *daf-2c(pe2722).* For both olfactory learning and taste learning, *daf-16; ist-1* double mutants were different from either single mutant, indicating that *daf-16* and *ist-1* function at least partly in parallel (Figure 3B, 5B).

### Food deprivation and insulin signaling regulate DAF-2 levels in AWC^ON^

In the ASER neurons that mediate taste avoidance learning, mVenus-tagged DAF-2a protein is restricted to the cell body, while mVenus-tagged DAF-2c protein is present in the cell body, axons, and synaptic regions (Ohno, et al., 2014). We observed similar subcellular localization patterns of mVenus-tagged DAF-2a and DAF-2c in AWC^ON^. DAF-2a was present in the AWC^ON^ dendrite and cell body, with a somatic pattern suggesting nuclear exclusion, but was undetectable in the AWC^ON^ axon (Figure 6A-C), whereas DAF-2c was present in the dendrite, cell body and axon of AWC^ON^ (Figure 6D-F).

**Figure 6.**
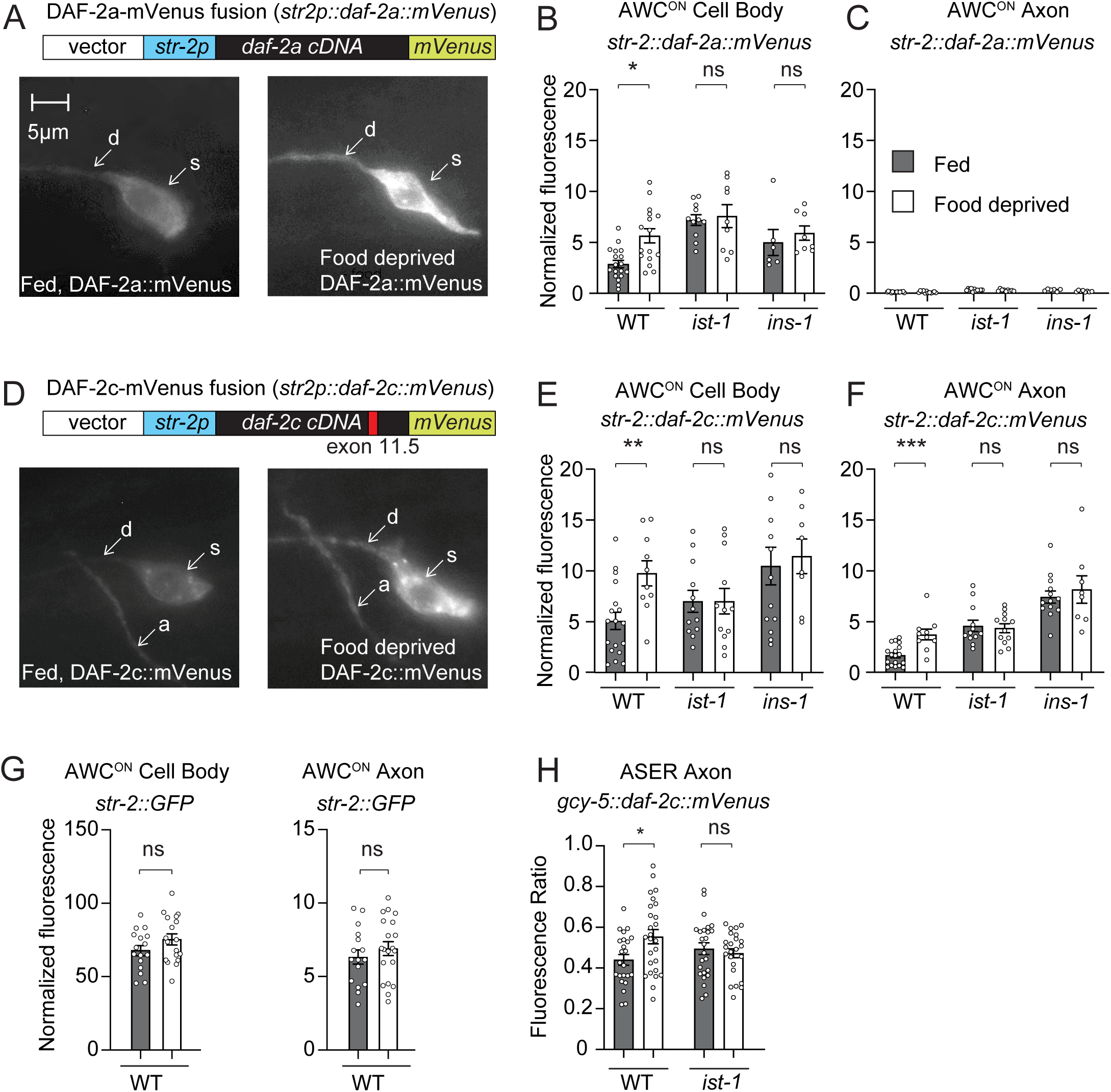
DAF-2 is regulated by nutritional state, *ist-1,* and *ins-1*. A) and D) Expression of DAF-2a::mVenus and DAF-2c::mVenus in AWC^ON^ neurons of fed and food-deprived animals. Top, diagrams of expression constructs. Bottom, representative maximal intensity projection of AWC^ON^ in well-fed or 90 min-food-deprived animals (d: dendrite, a: axon, s: cell soma). Note axonal expression of DAF-2c::mVenus and axonal exclusion of DAF-2a::mVenus. B) and C) DAF-2a::mVenus, E) and F) DAF-2c::mVenus fluorescence in AWC^ON^ cell bodies or axons, quantified either immediately after removal from food or 90 minutes after food deprivation on NGM agar plates. Expression of DAF-2a::mVenus (in the cell body) and DAF-2c::mVenus (in the cell body and axon) is increased by food deprivation in wild-type animals but not in *ins-1* or *ist-1* mutants. G) Quantification of *str-2::*GFP signals in wild-type animals either immediately after removal from food or 90 minutes after food deprivation on NGM agar plates. Food deprivation does not increase GFP expression in the AWC^ON^ axon or cell body. H) Expression of DAF-2c::mVenus in ASER neurons, quantified either immediately after removal from food or 90 minutes after food deprivation on NGM agar plates. Expression of DAF-2c::mVenus is increased in wild-type animals but not in *ist-1* mutants. In B,C,E,F, and G, Normalized fluorescence represents background-subtracted fluorescence of animals taken under identical imaging conditions across panels. Error bars indicate S.E.M. *, values differ at p ≤ 0.05. ns, values are not significantly different. In H, ASER DAF-2c::mVenus fluorescence was expressed as a ratio relative to a co-expressed mCherry transgene. Statistical tests are described in Table S2.

The levels of tagged DAF-2a and DAF-2c protein in AWC^ON^ increased after 90 minutes of food deprivation, the unconditioned stimulus used for aversive conditioning, but their subcellular localization patterns were unchanged (Figure 6A-F). In control experiments, expression of GFP under the same AWC^ON^ -selective *str-2* promoter was not affected by food deprivation, suggesting that DAF-2 translation, trafficking, or stability is altered post-transcriptionally (Figure 6G).

In mammals, insulin signaling results in internalization of insulin receptors, which may subsequently return to the cell surface or be degraded (Chen, et al., 2019). With that in mind, we asked whether insulin signaling itself might contribute to the observed changes in DAF-2 protein expression. Indeed, food deprivation did not affect AWC^ON^ DAF-2 levels in *ist-1* or *ins-1* mutants (Figure 6B-C, Figure 6E-F). Since *ist-1* has a minor role in salt learning, we asked whether *ist-1* also plays a role in the localization of DAF-2c in the salt-sensing ASER neuron. The levels of mVenus-tagged DAF-2c in the ASER axon increased after 90 minutes of food deprivation in wild-type animals, but not in *ist-1* mutants (Figure 6H). These results suggest that food regulates DAF-2 protein in sensory neurons via *ins-1* and *ist-1*.

### Aversive learning suppresses odor-dependent glutamate release from AWC^ON^

To ask how *ist-1-*dependent insulin signaling affects AWC^ON^ responses to odor in naïve and trained animals, we employed the genetically-encoded calcium indicator GCaMP5. AWC^ON^ calcium responses to butanone pulses were normal in *ist-1* mutants across a 100,000-fold range of butanone concentrations, consistent with their normal chemotaxis (Figure 7A-B). Sustained exposure to high butanone, with or without food, causes a ten-fold reduction in AWC^ON^ butanone sensitivity across its dynamic range that persists for at least 30 minutes (Cho, et al., 2016). This nonassociative AWC^ON^ desensitization was similar in wild-type and *ist-1* animals (Figure 7B). We conclude that the aversive learning defect in *ist-1* mutants arises downstream of the AWC^ON^ calcium signal, as previously proposed for *ins-1* and *age-1* insulin/IGF pathway genes (Cho, et al., 2016).

**Figure 7.**
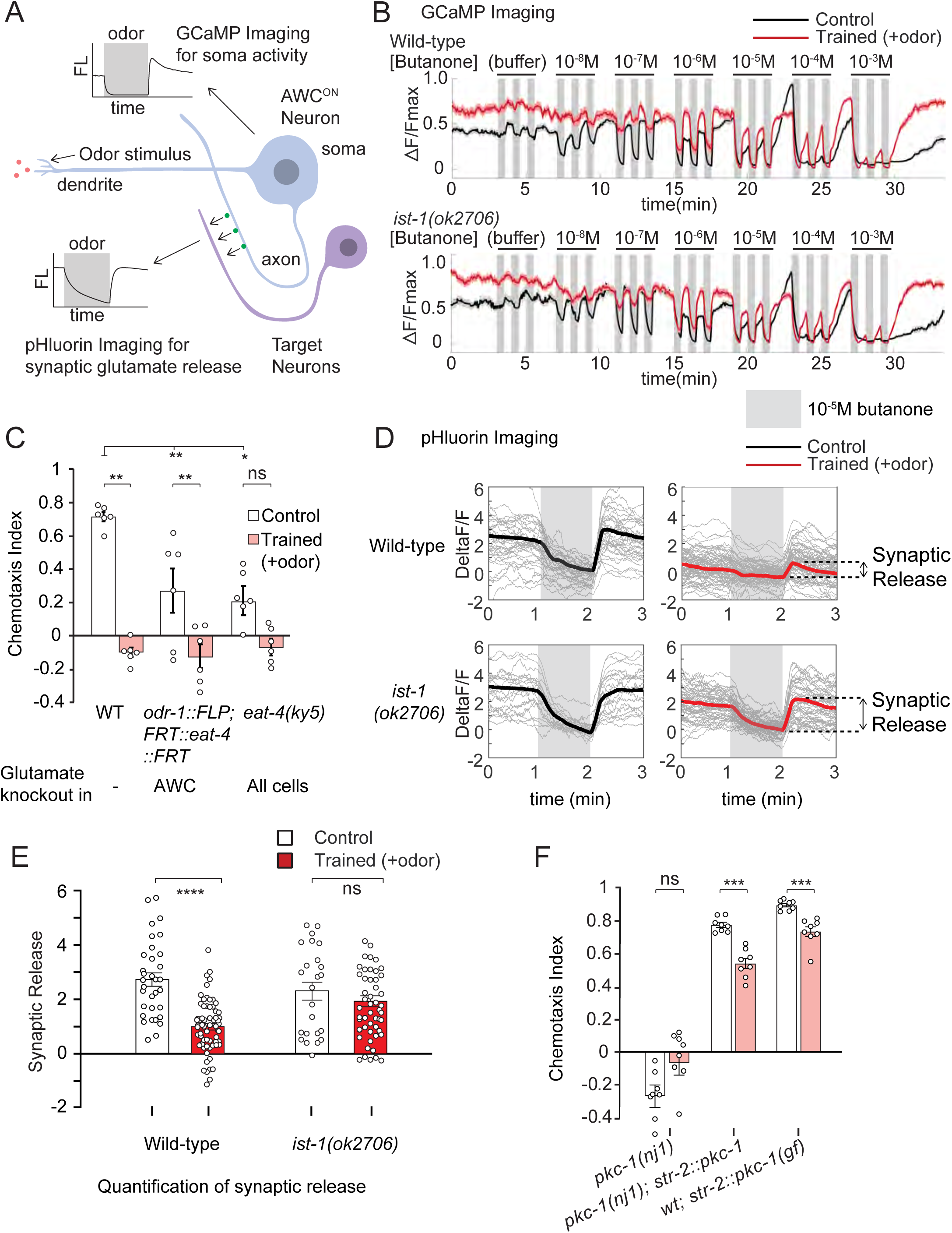
Aversive learning suppresses synaptic odor response in AWC^ON^. A) Diagram of AWC^ON^ neuron illustrating sites of odor detection in the cilia, the site of calcium imaging in the cell body, and the sites of synaptic EAT-4::pHluorin imaging in the axon. AWC^ON^ responds to odor removal with calcium increases that trigger synaptic vesicle release. B) Desensitization of the AWC^ON^ calcium response. Animals were exposed to 10 mM butanone for 90 minutes, or incubated in buffer for 90 minutes, followed by monitoring AWC^ON^ GCaMP5 responses to different odor concentrations. After butanone exposure (red lines), the AWC^ON^ dose-response curve is shifted ∼10-fold to the right compared to control animals (black lines) across the full range of odor concentrations. *ist-1* mutants showed similar desensitization to wild-type animals. Grey stripes indicate times when butanone stimulus was present (30s pulses, separated by 30s or 90s in buffer). Solid lines represent average data of multiple trials (WT control: n=64 worms from 7 independent experiments, WT trained: n=47 worms from 5 independent experiments, *ist-1(ok2706)* control: n=65 worms from 7 independent experiments, *ist-1(ok2706)*: n=55 worms from 5 independent experiments). Shading represents S.E.M. C) Chemotaxis and aversive learning in wild type, *eat-4* vesicular glutamate transporter mutants, and animals with AWC-selective knockout of *eat-4.* Animals were food-deprived (control, white bars) or food-deprived with 20 µl butanone odor (trained, pink bars) for 90 min and tested for chemotaxis to 1:1000 butanone. D) AWC^ON^ EAT-4-pHluorin responses to butanone pulses in wild type (top) and *ist-1* mutant (bottom) animals, in control (left, black lines) and trained (right, red lines) groups. Bold line shows average across multiple trials; thin lines show signal from each individual trial. Grey stripes indicate times when butanone stimulus was present (10^5^ M, 1 min pulses, separated by 2 min). E) Quantification of AWC^ON^ EAT-4-pHluorin signals from D). To calculate the change in synaptic release, the fluorescence signal during odor stimulation (5-10 s before odor removal) was subtracted from the post-stimulation fluorescence level (5-10 s after odor removal). (WT control: n=11 worms with 2-3 trials each, WT trained: n=22 worms with 2-3 trials each, *ist-1(ok2706)* control: n=10 worms with 2-3 trials each, *ist-1(ok2706)*: n=18 worms with 2-3 trials each). F) Chemotaxis and aversive learning in *pkc-1* mutants (see Fig 3A), *pkc-1* mutants with *pkc-1* rescued in AWC^ON^, and wild-type animals expressing *pkc-1(gf)* in AWC^ON^. Animals were trained and tested as above. Error bars indicate S.E.M. *, values differ at p ≤ 0.05. **, values differ at p ≤ 0.01. ***, values differ at p ≤ 0.001. ****, values differ at p ≤ 0.0001. ns, values are not significantly different. Statistical tests are described in Table S2.

The AWC^ON^ neuron releases the classical neurotransmitter glutamate and a number of neuropeptides (Chalasani, et al., 2010; Chalasani, et al., 2007). An AWC-selective knockout of *eat-4*, the vesicular glutamate transporter, was generated using FLP::FRT recombination targeted to the endogenous *eat-4* locus (López-Cruz, et al., 2019). AWC knockout of *eat-4* resulted in a butanone chemotaxis defect comparable to that of the *eat-4* null mutant (Figure 7C). These animals had a slight reduction in chemotaxis after pairing butanone and food deprivation, as did *eat-4* null mutants. Thus, AWC glutamate release is important for butanone chemotaxis, but likely functions together with additional AWC neurotransmitters.

To investigate the synaptic properties of AWC^ON^, we used a reporter strain in which the EAT-4 transporter was fused to pHluorin, a pH-sensitive version of GFP whose fluorescence increases when acidic glutamatergic synaptic vesicles fuse with the plasma membrane (Ventimiglia & Bargmann, 2017). Naïve wild-type animals and *ist-1(ok2706)* mutants had similar AWC^ON^ responses to butanone addition and removal using this synaptic reporter (Figure 7D, E). After pairing butanone with food deprivation, EAT-4::pHluorin responses to odor pulses were reduced in wild-type AWC^ON^ neurons for at least 30 minutes, consistent with a dampened synaptic response to odor (Figure 7D, E). Importantly, this synaptic effect was observed at butanone concentrations that elicited strong GCaMP responses in both naïve and trained animals (Figure 7B). By contrast, AWC^ON^ neurons in *ist-1* mutants retained substantial odor-regulated glutamate release after training, providing a synaptic correlate of their defect in aversive learning (Figure 7D, E). These results suggest that aversive learning reduces odor-regulated glutamate release from AWC^ON^ via *ist-1*-dependent insulin/IGF signaling.

To further explore the relationship between synaptic transmission and aversive learning, we examined PKC-1, a presynaptic protein kinase C homolog that stimulates glutamate release from AWC^ON^ neurons (Figure 3A) (Ventimiglia & Bargmann, 2017) and neuropeptide secretion (Sieburth, et al., 2007). PKC-1 is activated by Gq-PLC signaling; in AWC^ON^, *pkc-1* and Gq are key effectors of appetitive learning after pairing butanone with food (Arey, et al., 2018). *pkc-1* mutants were repelled by butanone, and only slightly affected by pairing butanone with food deprivation (Figure 7F). Their chemotaxis defect was rescued by expressing *pkc-1* in AWC^ON^, and interestingly, the rescued strain was defective in aversive learning (Figure 7F). The learning defect might be due to *pkc-1* overexpression in AWC^ON^ or the absence of *pkc-1* in another cell type, so we next examined a strain in which a *pkc-1(gf)* cDNA was specifically expressed in AWC^ON^ in wild-type animals. The AWC^ON^::*pkc-1(gf)* strain was highly defective in aversive learning, consistent with a role for AWC^ON^ presynaptic regulation in learning and memory (Figure 7F).

## DISCUSSION

These results provide three new insights into insulin/IGF signaling in olfactory learning. First, aversive learning in AWC^ON^ engages a cell type-specific insulin-*daf-2* signaling pathway that includes the adaptor protein IST-1. Second, insulin/IGF signaling causes DAF-2c insulin receptor levels to rise during food deprivation, suggesting an acute function that sets the stage for learning. Third, aversive training inhibits odor-regulated glutamate release from AWC^ON^ via the insulin/IGF signaling pathway. Through cell-specific inactivation of the endogenous *daf-2* locus, and through the analysis of *ist-1* mutants that have restricted defects in insulin/IGF signaling, we were able to localize *ist-1* and *daf-2* functions in aversive learning to AWC^ON^ sensory neurons. These genetic tools allowed us to avoid systemic developmental and metabolic effects of this critical endocrine pathway.

### Aversive learning uses a cell type-specific insulin/IGF signaling pathway

The insulin-related molecule encoded by *ins-1* and the insulin receptor encoded by *daf-2* are both required for aversive learning. By contrast, *ins-1* and *daf-2* have opposite effects on dauer formation (Kodama, et al., 2006). *C. elegans* has 40 genes that encode insulin-like peptides, and the *daf-2* gene encodes at least seven different protein isoforms with different functions (Ohno, et al., 2014; Martinez, et al., 2020). Structural studies of mammalian insulin and IGF-1 receptors predict that the DAF-2c-specific exon is adjacent to the primary ligand-receptor interface, and therefore could alter interactions between DAF-2 and insulin ligands (Gutmann, et al., 2019; Li, et al., 2019; Scapin, et al., 2018). We speculate that INS-1 activates DAF-2c, the isoform implicated in aversive learning, but inhibits the DAF-2a isoform used in dauer development.

In alternative models, INS-1 might not act directly on AWC^ON^, but indirectly via other cells. Insulin-to-insulin relays have been observed in several settings in *C. elegans*: for example, in pathogen-related learning, INS-7 from the URX sensory neuron antagonizes the insulin/IGF receptor DAF-2 in RIA interneurons, to prevent learning, while INS-6 secreted from the ASI sensory neurons suppresses INS-7 production from URX neurons, thereby disinhibiting RIA and allowing learning (Chen, et al., 2013). Several other neuronal insulins participate in long-term activity-dependent regulation that results in compensation across circuits (Yeon, et al., 2021). Results like these underscore the value of the cell type-specific *daf-2* knockout used here, which demonstrates a direct requirement for *daf-2* in AWC^ON^, although it leaves open the possibility that *daf-2* acts in other cell types as well.

Insulin and insulin-like signaling are usually associated with nutrients and growth in both vertebrates and invertebrates. However, the aversive learning paradigms for odor and salt in *C. elegans* require *ins-1, daf-2c,* and insulin signaling in the opposite context, food deprivation. *ins-1* signaling from the intestine drives food deprivation-induced thermosensory plasticity, consistent with a role for *ins-1* in encoding the low-food context (Takeishi, et al., 2020). Moreover, *ins-1* expression is upregulated by starvation both in the intestine and in the AIA neurons that regulate AWC plasticity (Takeishi, et al., 2020). We observed that DAF-2 levels in AWC^ON^ increase upon food deprivation through a post-transcriptional mechanism that requires *ist-1* and *ins-1*. These results suggest that acute insulin signaling during food deprivation results in altered translation, stabilization, or trafficking of DAF-2 receptors, and sets the stage for learning. There is precedent for such post-transcriptional regulation in mammals, where insulin/IGF receptors are internalized and may be degraded after ligand binding (Chen, et al., 2019), but IRS (IST-1) can promote retention of IGF-IR complexes on the plasma membrane instead of degradation by blocking an AP2 binding site (Yoneyama, et al., 2018).

IRS proteins are required for insulin/IGF signaling in mammals and in *Drosophila,* but were not previously thought to be important in *C. elegans* (Wolkow, et al., 2002). We found that *ist-1* has a prominent role in aversive olfactory learning. Expression of *ist-1* is highly enriched in AWC, ASE, AFD, and BAG sensory neurons, all of which show robust plasticity upon food deprivation. One possibility is that *ist-1* preferentially contributes to neuronal and behavioral *daf-2* signaling, and less to endocrine and developmental *daf-2* signaling in *C. elegans*; a second possibility is that IST-1 potentiates INS-1/DAF-2c interactions through inside-out signaling. Olfactory learning also has a strict requirement for the AKT-related kinase *akt-1,* whereas in development *akt-1* is largely redundant with *akt-2. akt-1* expression is enriched in many of the same neurons that express *ist-1,* including AWC, ASE, and BAG (Taylor, et al., 2021). Whether *akt-1* and *akt-2* differ only in their expression, or also in function, remains to be determined. Mammalian Akt1, Akt2, and Akt3 have different cellular and subcellular distributions (Santi & Lee, 2010), and distinct roles in synaptic plasticity in the hippocampus (Levenga, et al., 2017).

The analysis of double mutants suggests that insulin/IGF receptor signaling has two targets in AWC^ON^: one that depends on the *daf-16* transcriptional regulatory pathway, and one that does not. Similarly, in salt chemotaxis, insulin signaling occurs through one pathway that requires *daf-2a* and *daf-16,* and a second pathway that is *daf-2c-*dependent and *daf-16-*independent (Nagashima, et al., 2019). Using temperature-sensitive mutations in *daf-2,* Lin et al. (2010) showed that insulin/IGF signaling is required both for the formation and for the retrieval of an aversive olfactory memory. It is possible that these temporally distinct functions are associated with molecularly distinct insulin/IGF signaling pathways.

### Aversive learning suppresses presynaptic glutamate release from AWC^ON^

Odor responses in AWC^ON^ are regulated by olfactory experience and/or insulin/IGF signaling across multiple timescales. Sustained butanone exposure with food results in non-associative desensitization of AWC^ON^ calcium responses to butanone, but this is insufficient for aversive learning (Cho, et al., 2016). Insulin/IGF signaling transiently suppresses AWC^ON^ calcium responses through a rapid feedback loop, but this functional plasticity is short-lived compared to aversive learning (Cho, et al., 2016; Chalasani, et al., 2010). Here, we show that sustained pairing of butanone odor with food deprivation results in suppression of odor-regulated glutamate release from AWC^ON^. This presynaptic depression requires association of food deprivation with odor, as expected for aversive learning, and is retained for at least 30 minutes, consistent with the hour-long chemotaxis assay. As glutamate release from AWC is required for efficient chemotaxis and the odor-regulated turning behavior used in chemotaxis, this functional change is likely to contribute to behavioral changes after learning (Tsunozaki, et al., 2008; Chalasani, et al., 2007). A similar mechanism has been proposed to play a role in aversive taste learning, where calcium responses to salt in ASER sensory neurons are preserved but synaptic pHluorin signals are decreased after training (Oda, et al., 2011). The detailed mechanism by which insulin signaling suppresses glutamate release remains to be determined, and might involve AGE-1 mobilization of lipids that regulate synaptic proteins such as protein kinase C (PKC-1) (Ohno et al., 2014), AKT-1-dependent protein phosphorylation, or regulation of gene expression.

How is food deprivation associated with odor cues? Olfactory signal transduction in AWC^ON^ is mediated by cGMP, and one site of odor-food convergence is the cGMP-dependent protein kinase EGL-4. When butanone is present for an hour or more, and DAF-2/AGE-1 signaling is active, EGL-4 translocates from the AWC^ON^ cytoplasm to the nucleus (Cho, et al., 2016; Lee, et al., 2010; O’Halloran, et al., 2009; Torayama, et al., 2007; L’Etoile, et al., 2002). Like INS-1, EGL-4 is required for aversive but not appetitive learning (Cho, et al., 2016; Torayama, et al., 2007). The sustained coincidence of EGL-4 and insulin/IGF activity when odor is paired with food deprivation could represent a signal for associative learning. Proteomic studies indicate that EGL-4 is enriched in presynaptic regions (Artan, et al., 2021), where it could interact with DAF-2 signaling components. As mammalian EGL-4 homologs potentiate synaptic release in nociceptive neurons (Luo, et al., 2012), it is possible that depletion of presynaptic EGL-4 could contribute to synaptic depression in AWC^ON^, along with EGL-4-dependent transcriptional changes in the nucleus.

Olfactory neurons in all animals consist of many classes with distinct odor responses. Recent results have shown that mouse olfactory sensory neurons regulate dozens of activity-sensitive genes within a few hours in a novel olfactory environment, in a cell type-specific fashion, resulting in altered odor responses (Tsukahara, et al., 2021). Food deprivation is a powerful motivating stimulus in all animals (Smith & Grueter, 2021), and insulin receptors are prominently expressed in the rodent olfactory bulb (Kuboki, et al., 2021; Aimé, et al., 2012). It will be interesting to ask whether insulin signaling can modify mammalian olfactory neurons to provide associative input to an odor context. More broadly, insulin resistance in type-2 diabetes is associated with increased risk of Alzheimer’s disease, whose early symptoms include defects in olfactory processing (Talbot, et al., 2012). We speculate that exploring the relationship between insulin/IGF and learning in mammals could lead to a deeper understanding of these clinical findings.

## AUTHOR CONTRIBUTIONS

D.C. and C.I.B. designed experiments. D.C. conducted molecular, cellular, and behavioral analyses of olfactory learning. J.L. conducted dauer formation experiments. M.T. and Y.I. conducted salt learning experiments. D.C., M.B. and M.E. generated the CRISPR *daf-2* conditional knockout line. D.C. and C.I.B. analyzed and interpreted results and wrote the paper with input from all authors.

## ACKNOWLEDGEMENTS

We thank the *Caenorhabditis* Genetics Center (NIH P40 OD010440) for strains, Patrick McGrath for sequence analysis of *ist-1(ky1071),* Rebecca Butcher for synthetic ascarosides, Christine Cho, Alejandro Lopez, Elias Scheer, Navin Pokala, May Dobosiewicz, Aylesse Sordillo, and Donovan Ventimiglia, for advice and discussion, and Aylesse Sordillo, Elias Scheer, Yoav Printz, and Audrey Harnagel for comments on the manuscript. Du Cheng was supported by the Paul and Daisy Soros Fellowship for New Americans and by a Medical Scientist Training Program grant from the National Institute of General Medical Sciences of the NIH (T32GM007739) to the Weill Cornell/Rockefeller/Sloan Kettering Tri-Institutional MD-PhD Program. This work was supported by the Howard Hughes Medical Institute, of which Cori Bargmann was an investigator, and by the Chan Zuckerberg Initiative.

## KEY RESOURCES TABLE

**Table 1.**
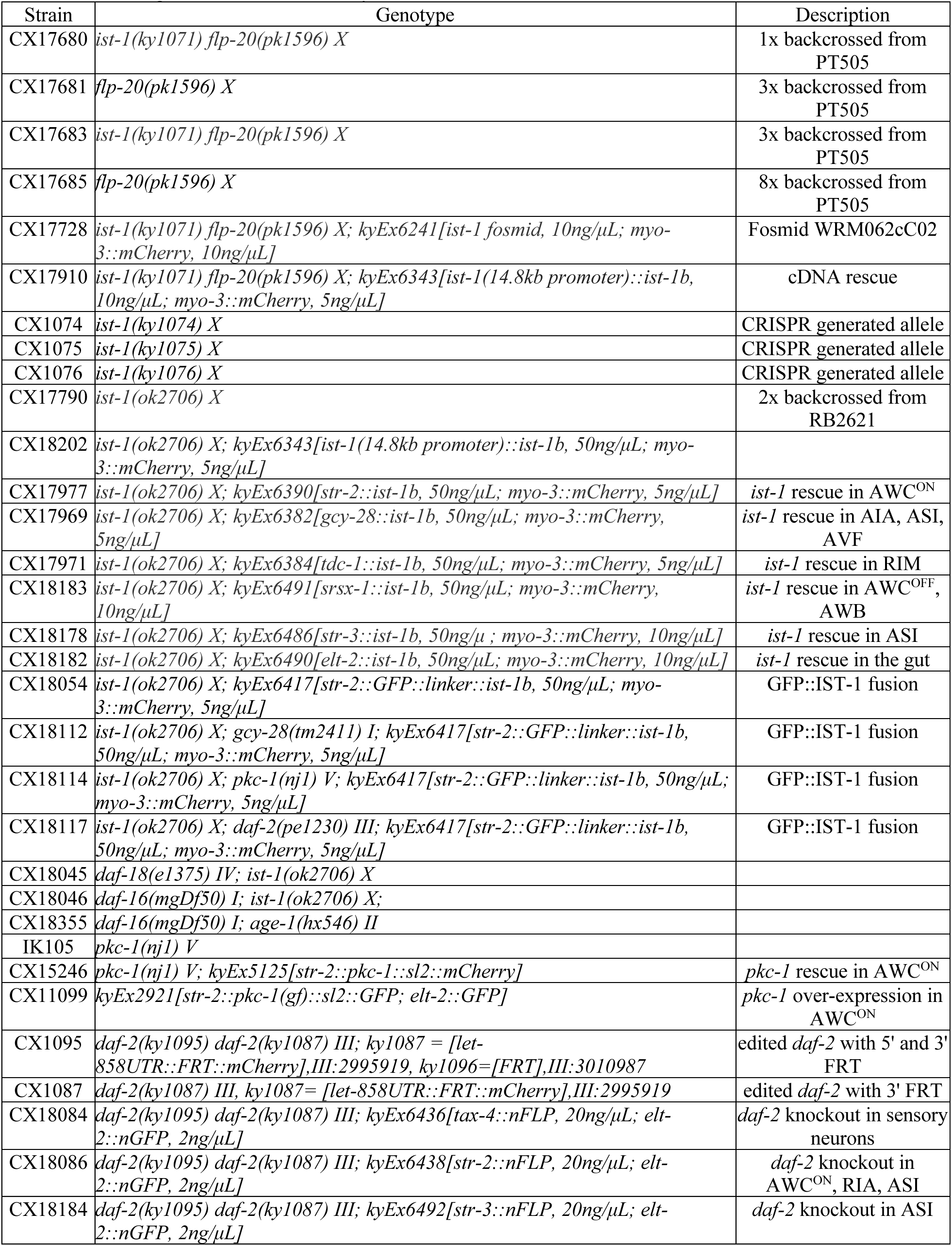
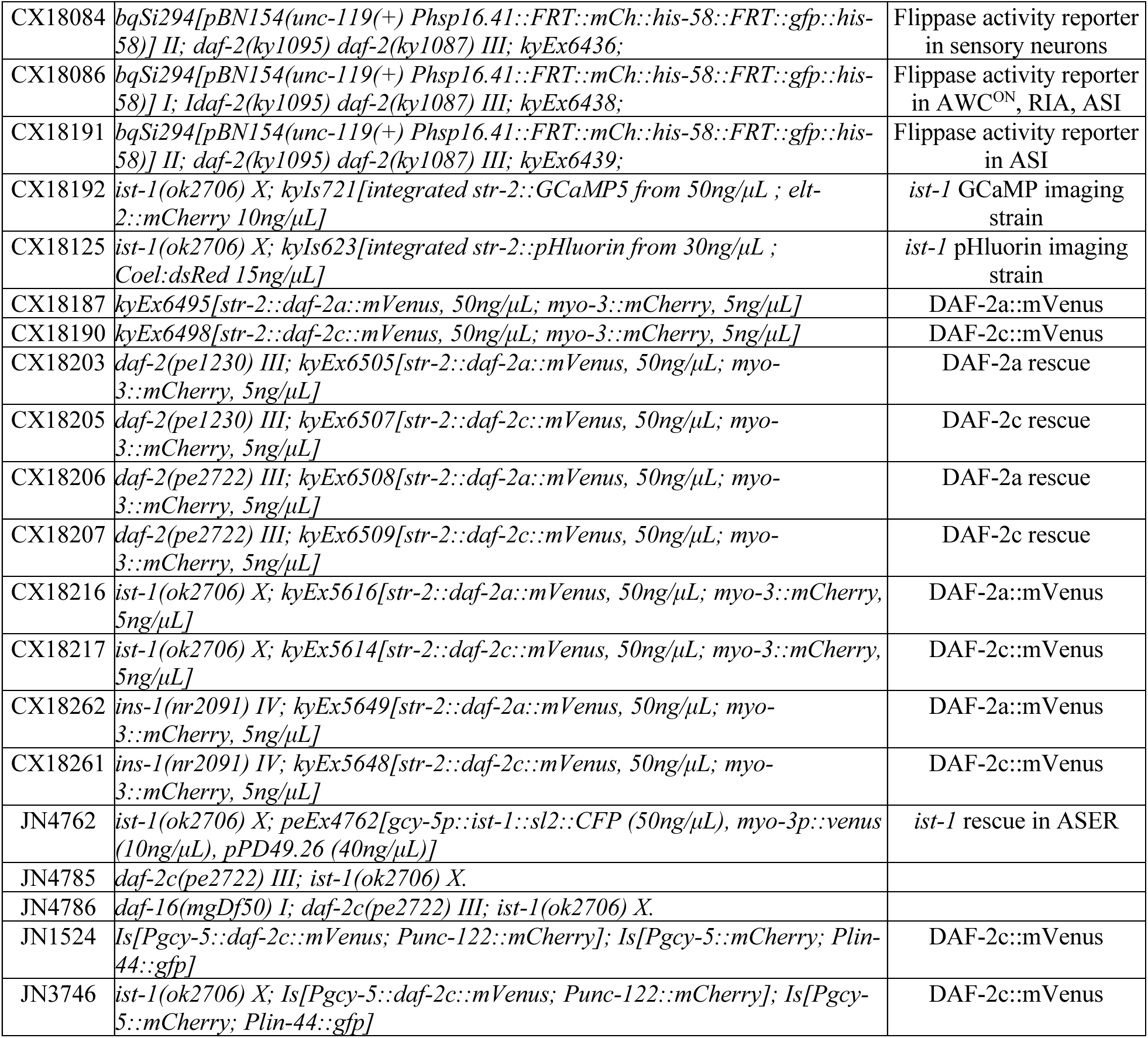
Strains generated in this study.

## CONTACT FOR REAGENT AND RESOURCES SHARING

Further information and requests for resources and reagents should be directed to and will be fulfilled by the Lead Contact, Dr. Cornelia Bargmann.

## EXPERIMENTAL MODEL AND SUBJECT DETAILS

### *C. elegans* strains

All new CRISPR-generated strains and transgenic strains are listed in Table 1.

### Nematode maintenance and molecular biology

All strains were maintained at 20°C on Nematode Growth Medium (NGM) plates, seeded with *E. coli* OP50 bacteria as a food source (Brenner, 1974). Wild-type animals were the genetically defined strain PD1074 derived from the Bristol strain N2 (Yoshimura, et al., 2019).

## METHOD DETAILS

### Chemotaxis assays

Animals were grown at 20°C on 10 cm round NGM plates spotted with 900 µl of *E. coli* OP50. Chemotaxis plates are square plates containing 12 ml of modified NGM agar (without peptone and cholesterol, 1.7% Granulated, purified agar (Agar-agar, Millipore), 50 mM NaCl, 25 mM KHPO_4_ (pH6), 1 mM CaCl_2_, 1 mM MgSO_4_). The chemotaxis plates are allowed to air-dry without lids for 15 minutes before use, or are stored at 4°C for up to 2 weeks before use. Fifteen minutes before the start of the assay, one 1 µl spot of NaN_3_ was spotted at the midline of each end of the plate. Adult animals were washed three times with NGM buffer (50 mM NaCl, 25 mM KHPO_4_ (pH6), 1 mM CaCl_2_, 1 mM MgSO_4_) with 15 min of total wash time. 50-100 animals were then placed at the center of the square plate spotted with either one 2 µl drop of butanone diluted to 1:10 or 1:1000 in ethanol, or an ethanol control. A piece of folded, absorbent paper (Kimwipe) was used to soak up the solution from the droplet before the lid was closed, freeing the animals from the droplet and beginning the assay. The assay was recorded in an imaging apparatus with Streampix software and a 6.6 MP PL-B781F CMOS camera (Pixelink) as described in (López-Cruz, et al., 2019) for 1 hour at 3 frames/second. The chemotaxis index was quantified at the last frame of the recording by counting animals that had left the origin and accumulated on the side closest to the odor (#Odor), on the side closest to the control (#Control), and in the intermediate region (#Middle). Animal positions were also tracked for further analysis. The formula used to calculate the chemotaxis index was: Chemotaxis index = (#Odor - #Control) / (#Odor + #Control + #Middle)

### Training for aversive learning

The training protocol for aversive learning is based on the assay developed by (Colbert & Bargmann, 1995) with modifications. One-day old adults were washed three times with NGM buffer and placed on conditioning plates consisting of NGM agar without peptone and cholesterol (same as chemotaxis plates). For the trained group, 20 µl of butanone were distributed on four small pieces of agar on the inside of the lid of the plate, while the control group did not have butanone on the lid. The plates were then sealed with parafilm for 90 minutes before animals were washed 2-3 times with NGM buffer and tested in chemotaxis assays according to the previous section.

### Training for desensitization and appetitive learning

The training protocol for desensitization and appetitive learning (or enhancement) is identical; the two assays are distinguished by test conditions (Torayama, et al., 2007). One-day old adults were washed three times with NGM buffer and placed on conditioning plates consisting of NGM agar seeded with 900 ul of OP50. For the trained group, 20 µl of butanone were distributed on four small pieces of agar on the inside of the lid of the plate, while the control did not have butanone on the lid. The plates were then sealed with parafilm and incubated for 90 minutes before animals were washed and tested in chemotaxis assays according to the section “**Chemotaxis assays**” with 1:1000 butanone for desensitization, and 1:10 butanone for appetitive learning.

### Salt concentration learning assay

Salt concentration learning assays were performed according to our previously reported procedure (Tomioka, et al., 2016). For conditioning on agar plates, adult worms were transferred to NGM plates with 25- or 100-mM NaCl in the absence or presence of a bacterial food (NA22) for 4–5 h. After conditioning, the worms were placed at the center of a test plate with an NaCl gradient and allowed to crawl for 45 min. A chemotaxis index was determined according to the following equation: NaCl Preference Index = (NA–NB) / (Nall–Nc), where NA and NB are the number of worms in the high- and low-salt areas, respectively, Nall is the total number of worms on a test plate, and Nc is the number of worms in the area around the starting position.

### Chemotaxis recordings and analyses

40-60 animals were placed on a square plate with modified NGM agar. A 1-hour movie was recorded at 3 frames per second with Streampix software and a 6.6 MP PL-B781F CMOS camera (Pixelink). Animals’ trajectories were extracted by a custom MatLab script. A long-reversal-omega turn (pirouette) event was defined as a reversal coupled with an omega turn, and identified as a sharp change in angular speed (≥75 degree/s), followed by a sharp reorientation (body-enclosing ellipse eccentricity ≤0.875, filtered by an angular speed threshold ≥60 degree/s), as previously described (Pokala, et al., 2014). Long-reversal-omega frequency was calculated by binning the events with respect to the angle between each worm’s heading and the direction of the odor source before the reorientation, at 30° intervals. The normalized long-reversal-omega frequency was calculated by binning the event/worm/minute with respect to angle between each worm’s heading and the direction of the odor source before the reorientation, at 90° intervals.

### Dauer formation assays

Dauer formation assays were performed using synthetic dauer-inducing pheromones with modifications to previously described methods (Neal, et al., 2013; Butcher *et al*., 2008). Assay plates were prepared 24 hours before each experiment by mixing 1 ml NGM-agar with 100 μl synthetic pheromone mixture (ascarosides C3 and C6, 73 nM final concentration each) or ethanol control. On the day of each experiment, 20 μl heat-killed OP50 (8 mg/ml in NGM buffer) was added and dried onto each plate. 14 adult animals were picked onto each assay plate and allowed to lay eggs for 3-4 hours before being removed. Plates were sealed with parafilm and incubated at 25°C with wet paper towels to prevent desiccation. After 72 hours of incubation, the proportion of dauers (including partial dauers) in each assay plate was counted based on their distinguishing characteristics: radial constriction, posture, gut refraction, locomotion, and cessation of pumping (Neal, et al., 2013).

### Calcium imaging in the arena chip

Multiple animals expressing GCaMP5 in AWC^ON^ were imaged simultaneously as described (Larsch, et al., 2013; Cho, et al., 2016) with slight modifications. PDMS chips with two arenas were made using a custom mold. The chips were flooded with NGM buffer containing 10 mM tetramisole hydrochloride, and 10-15 adult animals were loaded into each arena. Animals were trained in the chip for 90 minutes with 10 mM butanone (or NGM buffer for control groups) followed by a 15-minute wash with NGM buffer, then imaged on a Zeiss AxioObserver A1 microscope with a 2.5X objective and Metamorph software for synchronized image capture with pulsed illumination. A three-way valve was used to switch between buffer and odor flow into the chip, and a Hamilton valve was used to switch between different concentrations of odor. The stimulation protocol was three 30-second alternations between odor and buffer followed by 90 seconds of buffer, and then the sequence was repeated at 10-fold higher odor concentration for a total of six butanone concentrations, ranging from 10^-^8 M to 10^-3^ M, plus a buffer-to-buffer control. Fluorescence was measured using a custom ImageJ script (Larsch, et al., 2013). Calcium imaging data analysis was identical to that used in (Cho, et al., 2016) using custom Matlab scripts to analyze calcium imaging data generated by ImageJ tracking programs.

### pHluorin imaging in the single-animal trapped chip

The pHluorin imaging method is similar to that developed by (Ventimiglia & Bargmann, 2017). 1-day old adults expressing an AWC^ON^::EAT-4-pHluorin transgene were picked off food and trained for aversive learning. After training, animals were washed 3 times for a total of 15 minutes with NGM buffer and loaded into the imaging chip. Polydimethylsiloxane (PDMS) imaging chips were fabricated as described in (Chronis, et al., 2007) using a custom-designed silicon mold. PDMS chips were cured and holes were punched in fluid inlets, and chips were then bonded to glass cover slips. Animals were immobilized in the chip by adding 1 mM tetramisole hydrochloride (Sigma) to the worm holding chamber. A time stack of images was captured using a CoolSnap HQ Photometrics camera and a Zeiss Axioscope upright microscope with a 40X objective. Images were motion-corrected using the method developed by (Tseng, et al., 2011). Fluorescence measurements were extracted with custom, semi-automated tracking software in ImageJ script (Ventimiglia & Bargmann, 2017) that tracked axon position and extracted intensity measurements. pHluorin imaging analysis was similar to that used in (Ventimiglia & Bargmann, 2017), with slight modifications. Data was smoothed with a moving average of 50 frames before analysis and plotting.

### Quantitative analysis of DAF-2::mVenus subcellular localization

For AWC^ON^ imaging, one-day old adult animals were either picked directly off plates (fed) or moved to an unseeded NGM plate for 90 minutes for food deprivation. Animals were then mounted and anaesthetized on a 3% agar pad containing 10 mM sodium azide. Images were taken within 5-30 minutes after mounting. Fluorescent images of live animals were acquired using a Zeiss Axio Imager Z1 Apotome system with an AxioCamMR3 camera through an excitation filter of BP 500/20nm, with a beam splitter at 515 nm and an emission filter at BP 535/30 with a 20% attenuator setting on the light source. A Z-Stack of 30 images was taken at 1 μm spacing with a 63X/1.40 NA objective with 500 ms exposure time per image. DAF-2a or DAF-2c::mVenus fluorescence intensity was measured with ImageJ. An image containing the largest area of the AWC^ON^ cell body or axon based on mVenus expression was selected from Z-series for quantification. DAF-2::mVenus quantification was different from that described in (Ohno, et al., 2014), and more similar to that described in (Nagashima, et al., 2019). A rectangular region of interest (ROI) was drawn manually inside the area of the cell body or a piece of axon before using ImageJ to calculate the average intensity of fluorescence in the ROI. Average fluorescence intensity from at least five ROIs was used to quantify axon fluorescence of a given image stack. Background fluorescence intensity was averaged from at least five ROIs adjacent to the cell body or axon. The normalized fluorescence was calculated by subtracting the average fluorescence intensity of the background from the average fluorescence intensity of the cell body or axon.

For ASER imaging, fluorescence ratio was calculated based on the relative signal intensities of DAF-2c::mVenus and mCherry co-expressed in ASER.

### Identification of *ist-1(ky1071)* allele from genomic sequencing of *flp-20(pk1596)*

Genomic DNA was prepared from wild-type and four strains resulting from backcrossing PT505 *flp-20(pk1596)* to wild-type. The strain CX17680, a 2X backcross of PT505 *flp-20(pk1596),* and the strain CX17683, a 6X backcross of PT505 *flp-20(pk1596),* were both defective in butanone aversive learning. The strain CX17681, a 6X backcross of PT505 *flp-20(pk1596)* and CX17685, an 8X backcross of PT505 *flp-20(pk1596),* were both proficient in butanone aversive learning. Whole genome sequencing of these samples was performed by the Rockefeller Genomic Resource Center using an Illumina HiSeq 2500 System. Sequence reads were analyzed by Patrick McGrath (Georgia Tech) by comparing the sequence of the backcrossed strains to the wild-type strains using a custom program. A T->G mutation was found in the splicing motif for *ist-1* after the end of the 3^rd^ exon and verified using Sanger sequencing. The *ist-1(ky1071)* point mutation, which is predicted to disrupt a splice donor, is: ggaatttcagTATCAATGGCAGCAAGAGAGGAGAGAATACGCGAAAGTAGACAACAAGGGCAACTGAAAGAAAAAAGGCTGAAAAATCCAGATTCAAAAGGTGATACAAGTCACGATAAGCCGACCAGAGAAACATGGAAGCCGTTAGCCGTTGAAGACTTACCAAAGGATGAAGATCCGGATGAATTTGGAATTCCAGAAGTCTACAAATGTGGAAATTGTTTAGTAGGGTTTGCTCCAGTGg<u>t[g]</u> atgtttatttagaaattatgtgcacaattcaattttttgagcagttgtcgatcaaaaagaaaattctttcagatttatttttaattatgcataaaatcttcatattcc (site of point mutation underlined in red, mutation in square bracket, 3^rd^ exon in uppercase, introns in lowercase)

### CRISPR/Cas9-generated mutant strains

New mutations in *ist-1* were generated using a co-CRISPR protocol (Arribere, et al., 2014), with a template-guided dominant mutation on the *rol-6* gene to induce a roller phenotype to aid selection of the gene-edited progeny. Young hermaphrodites with only a few eggs in the gonad were injected with a mix of plasmids encoding Cas9, a gRNA targeting *rol-6*, a gRNA targeting the gene of interest (such as *ist-1*), and a ssDNA repair template that induces the *rol-6(su1006)* mutation. F1 animals carrying roller phenotypes were isolated onto individual plates, allowed to lay eggs, and then screened for a target mutation by Sanger sequencing. non-Roller F2 animals with frameshift mutations in *ist-1* were chosen to generate homozygous strains. The homozygous strains were then backcrossed twice to wild-type strains before being used to test behavior.

The sequence of the *ist-1(ky1074)* mutation (resulting in a frameshift in exon 2 affecting isoforms 1b and 1e) is: AAAACAGTGAGCTTCAGCGC<u>TGCCAACAATAATA</u>AACGGTGTCTGAGCTGATGTTGCCCGCCGAGCC (Deletion underlined).

The sequence of the *ist-1(ky1075)* mutation (resulting in a frameshift in the third shared exon, affecting all isoforms) is: GCTAATGTTTGTGACAC<u>TCAC</u>GGAACGATGTTTAGAACTTCACGAGTCGGAGAAATCCTATCGTGCG (Deletion underlined).

The sequence of the *ist-1(ky1076)* mutation (resulting in a frameshift in exon 2 affecting all isoforms) is: GCGCCAGGAAGAAACAGGAAGAAACTGTAAAGAAATgtaagtgtcgttctgttatggttgttaaatt (Deletion underlined).

### Generating a conditional *daf-2* allele

The flippase-FRT-modified *daf-2* strain, including an FRT site sequence, the *let-858* 3’UTR, and the mCherry coding sequence before the 3’ UTR of the *daf-2* gene, was made using a protocol developed by the Mello lab (Dokshin, et al., 2018). Briefly, *S. pyogenes* Cas9 protein and synthesized tracrRNA and crRNA were used to generate the cut required for recombination. The *let-858* 3’ UTR-FRT-mCherry insertion sequence was first assembled by standard cloning methods into a plasmid vector. Then oligos were used to generate the repair template by PCR: a pair of 55 nt 5’SP9 (TEG) modified oligos added 35 nt of homology to each side of the insertion site in the genome. The repair template was then heated and cooled with specific downsteps in the thermal cycler (95°C-2:00 min; 85°C-10 sec, 75°C-10 sec, 65°C-10 sec, 55°C-1:00 min, 45°C-30 sec, 35°C-10 sec, 25°C-10 sec, 4°C-O/N. Ramp down at 1°C/sec). Although the mechanism is unknown, the Mello lab found these heating-cooling steps are critical to achieve high knock-in efficiencies. The repair template, tracrRNA, crRNA and *rol-6(su1006)* plasmid co-injection marker were then injected into young hermaphrodite gonads, and mutants were selected as described above. (See section “**CRISPR/Cas9-generated mutant strains**”). The upstream FRT site was inserted in the middle of the *daf-2* gene such that excision would delete exons 10 through 17 which encode all intracellular domains including the kinase domain. We note that the *daf-2b* isoform, which encodes a secreted protein that antagonizes insulin signaling, might be expressed in a shortened form (exons 1-9, instead of its normal sequence from exons 1-11) (Martinez, et al., 2020). For the second site, we inserted a *let-858* 3’UTR-FRT-mCherry coding sequence between the stop codon and the 3’ UTR of *daf-2,* with the intention that mCherry should be expressed after successful knockout of *daf-2* in cells of interest. Due to oversight in design, the mCherry reporter could not be visualized after FRT recombination. Instead, the BN294 *bqSi294* strain, which carries an integrated transgene that expresses ubiquitous mCherry and a masked GFP, was used to report cell-specific flippase expression (Muñoz-Jiménez, et al., 2017). Upon flippase recombination between two FRT sites, mCherry is excised and GFP expression is unmasked. In the *str-2p::flippase bqSi294*strain, GFP was present in AWC^ON^ in 7/11 animals, in ASI in 4/11 animals, in URX in 4/11 animals, and less frequently in OLQ, CEP, AFD, ASK, RIC, pharyngeal muscle cells, and body wall muscles cells (1/11 to 3/11 animals each).

The sequence surrounding the 5’ insertion of the FRT site in *daf-2* (CX1095) conditional knockout strain is:

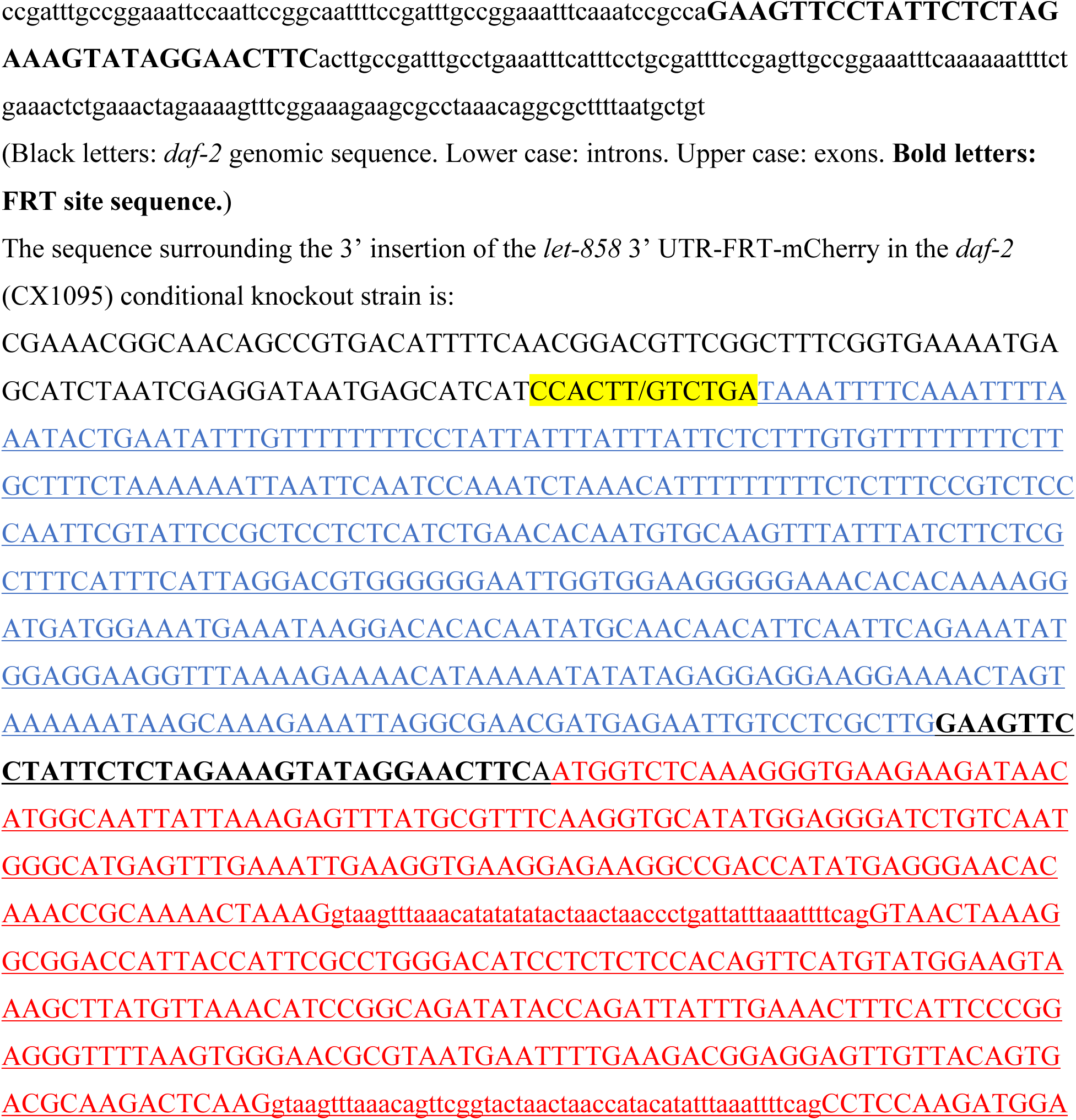
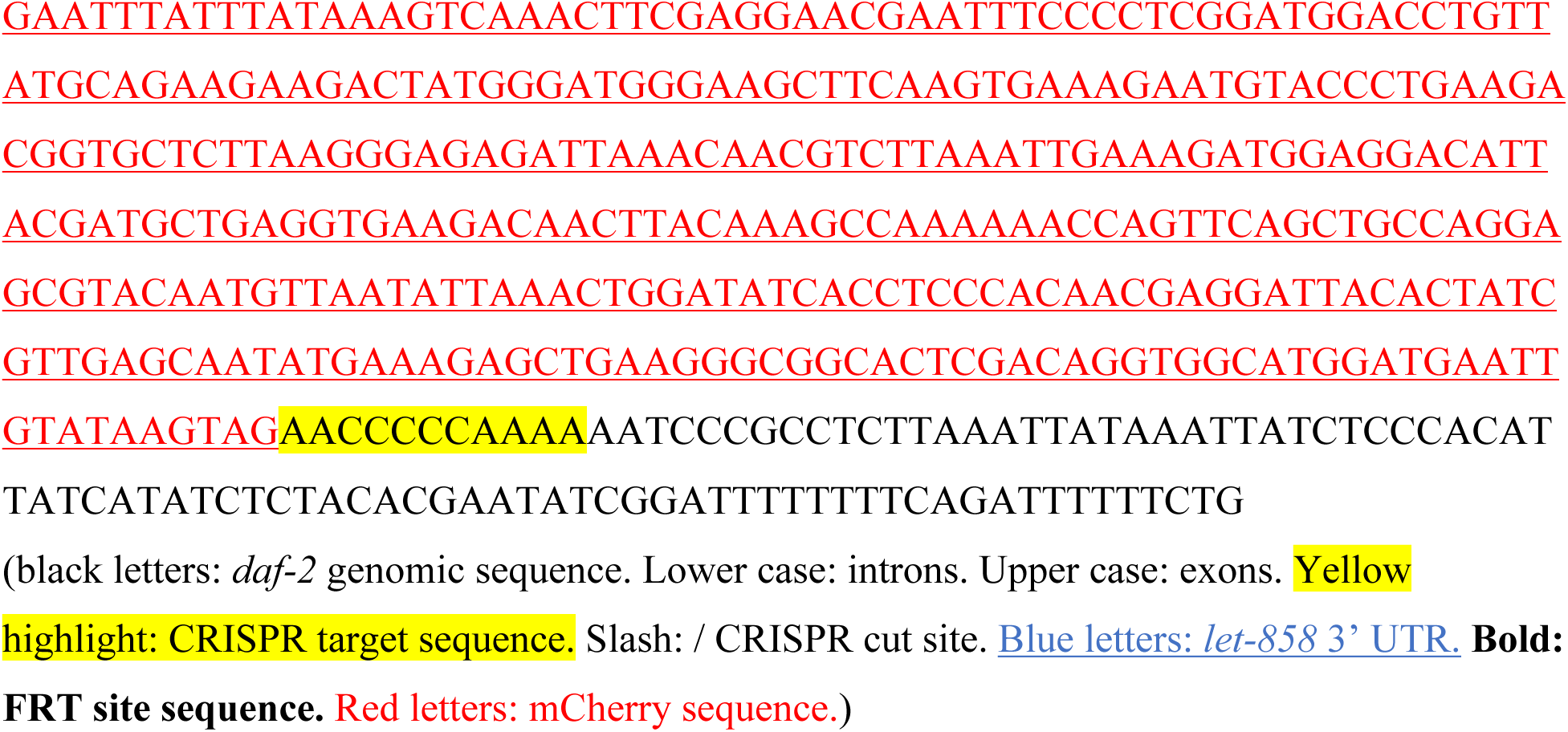

**Table 2.**
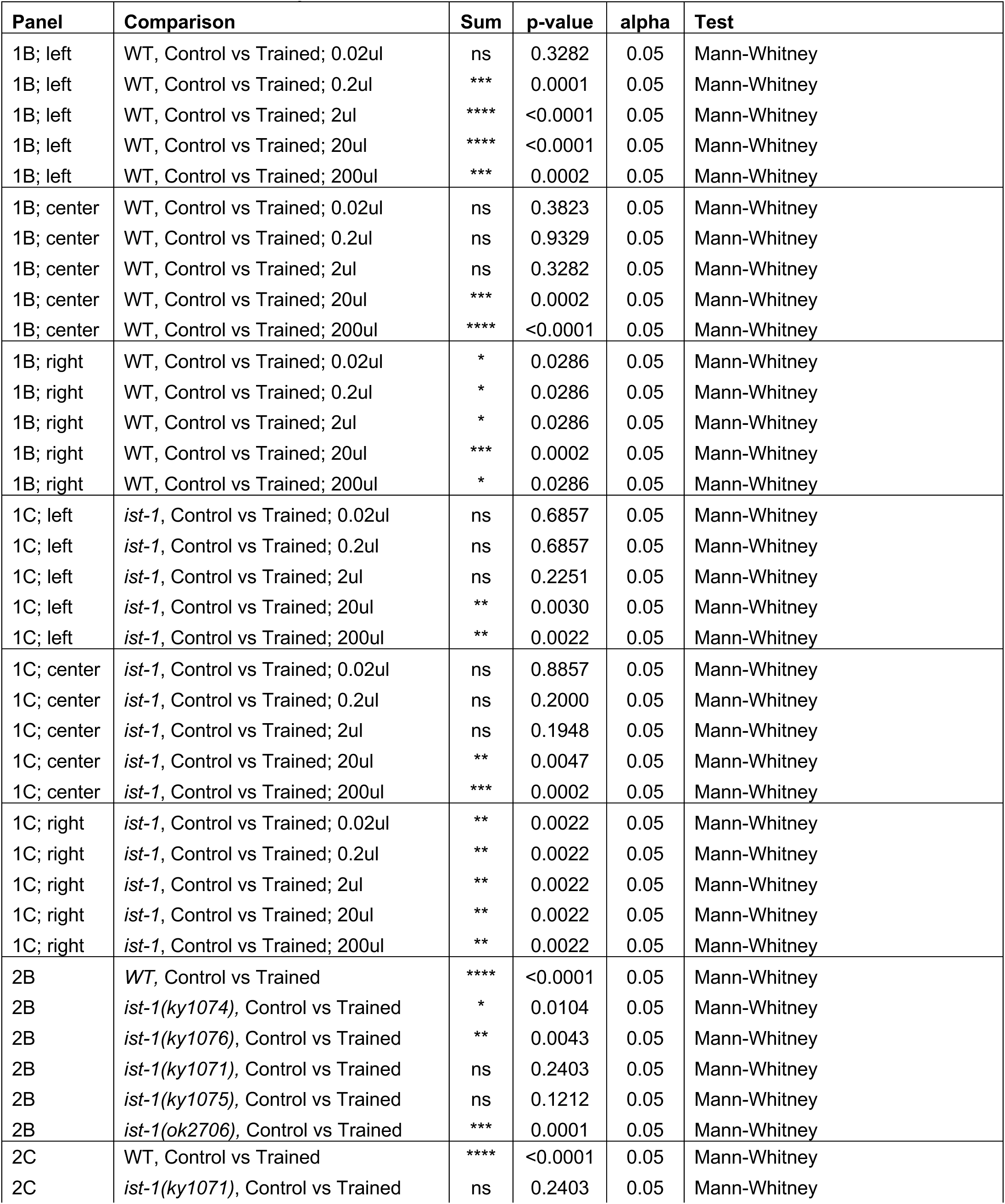
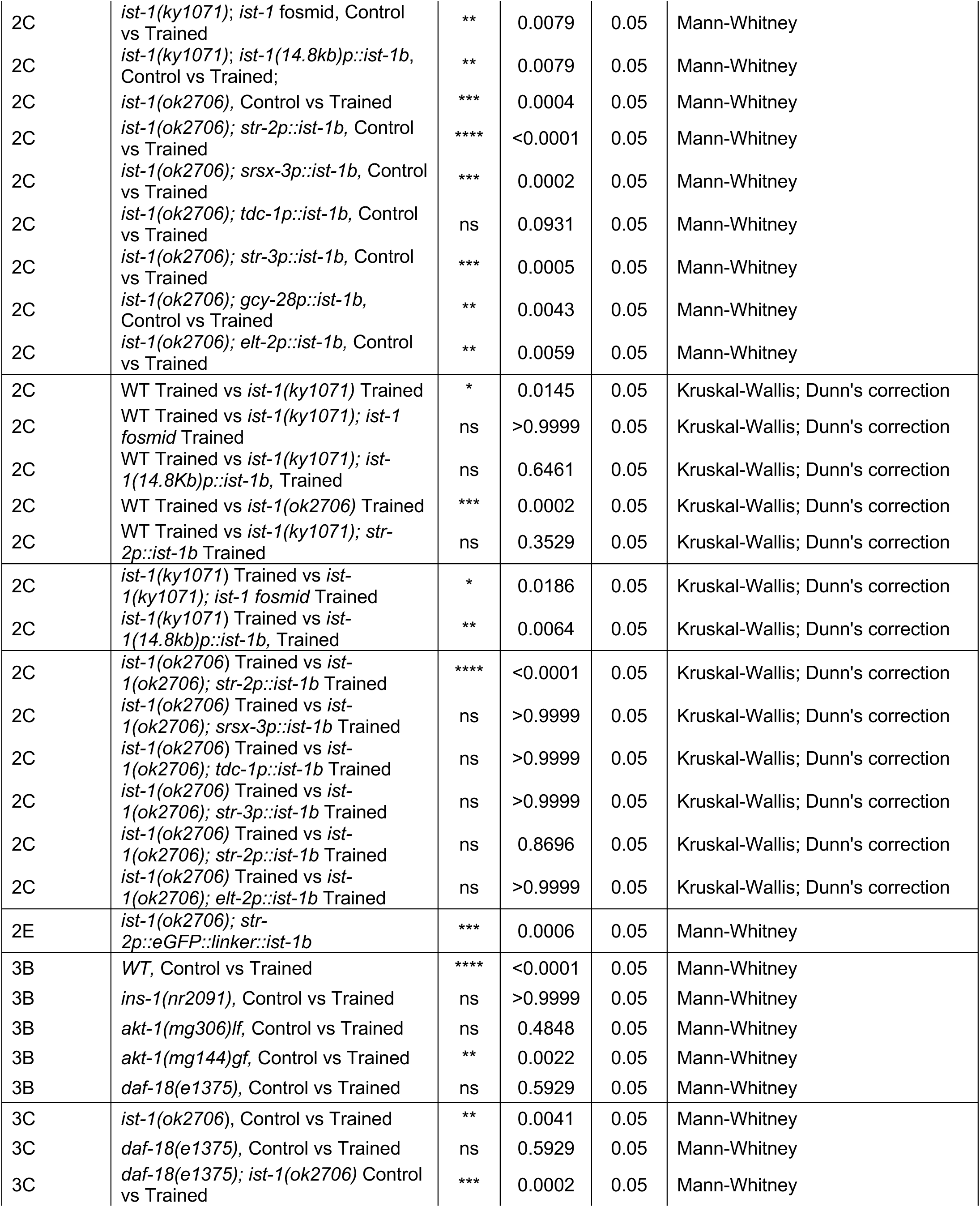
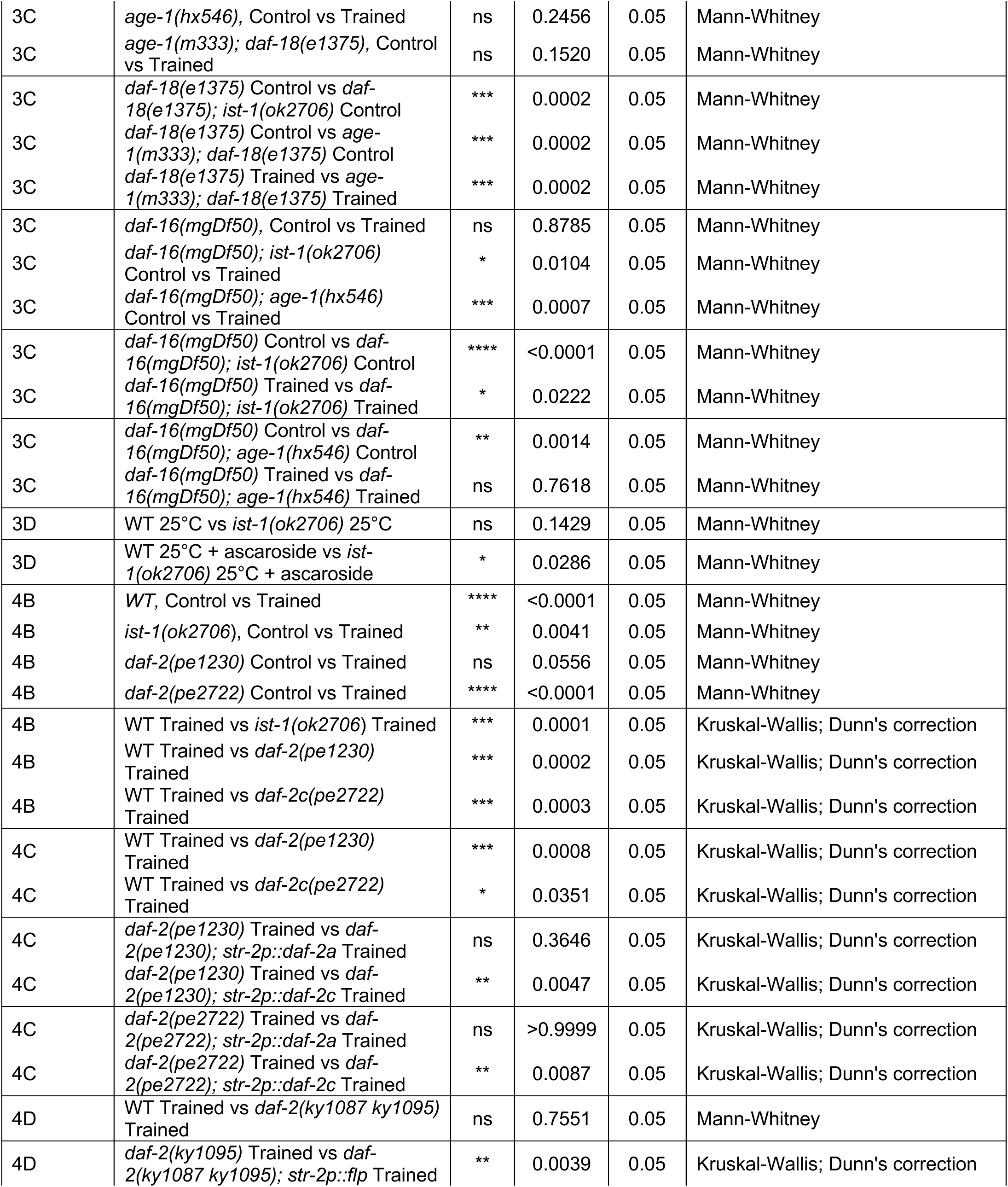
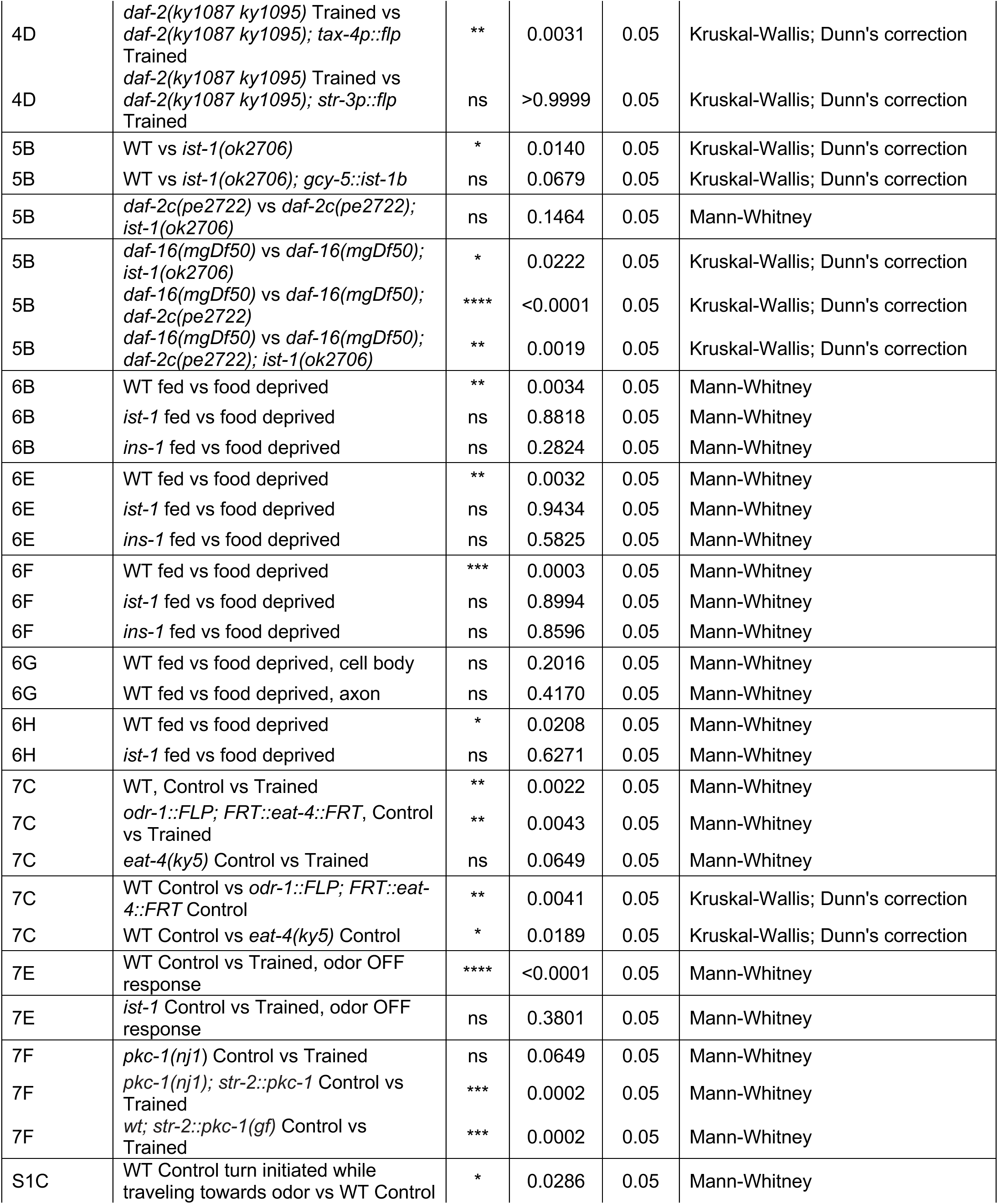
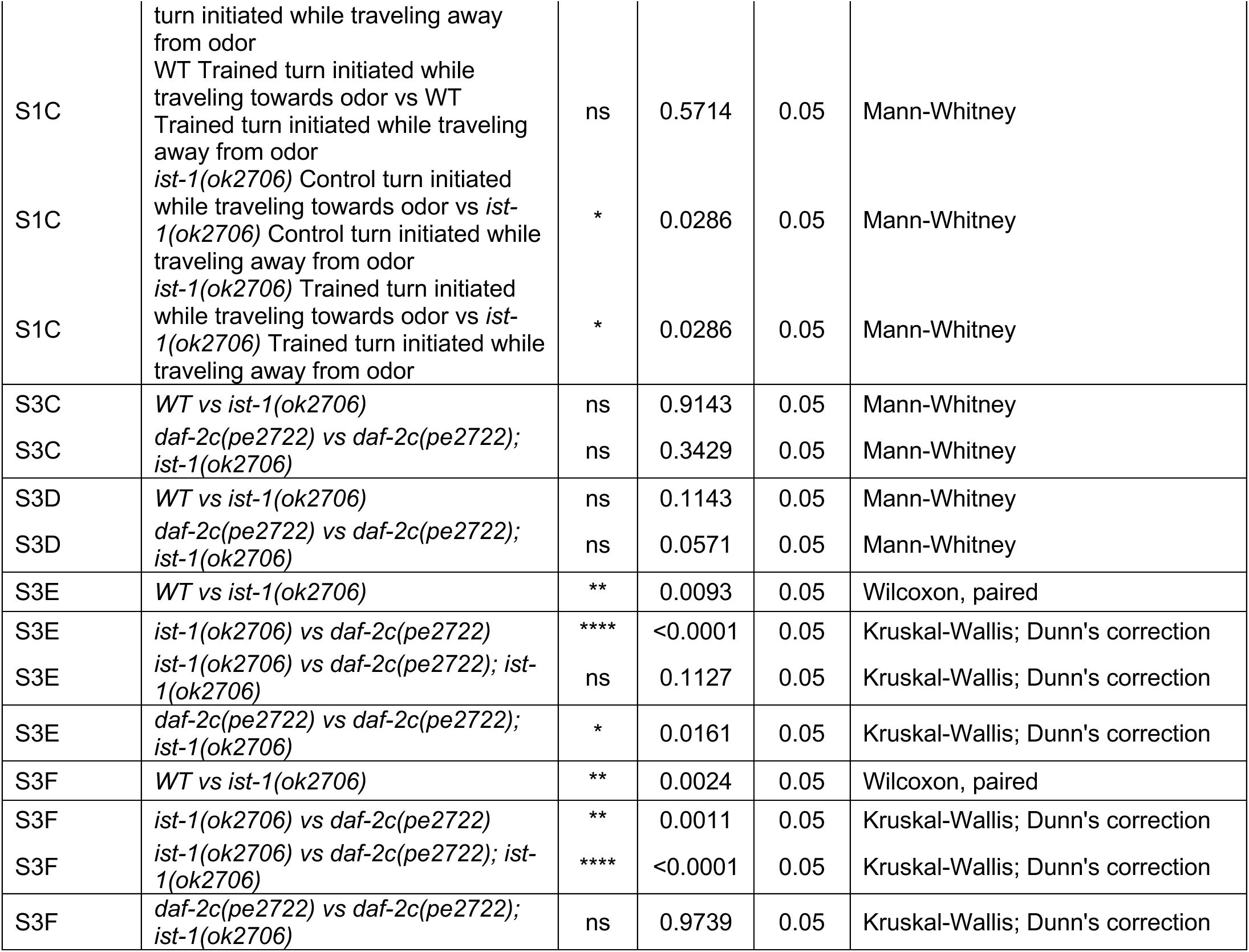
Statistical Analyses.

## DATA AND SOFTWARE AVALIABILITY

All datasets and scripts generated during the current study will be made available on Mendeley Data and Github, respectively.

## SUPPLEMENTARY FIGURE LEGENDS

**Figure S1.**
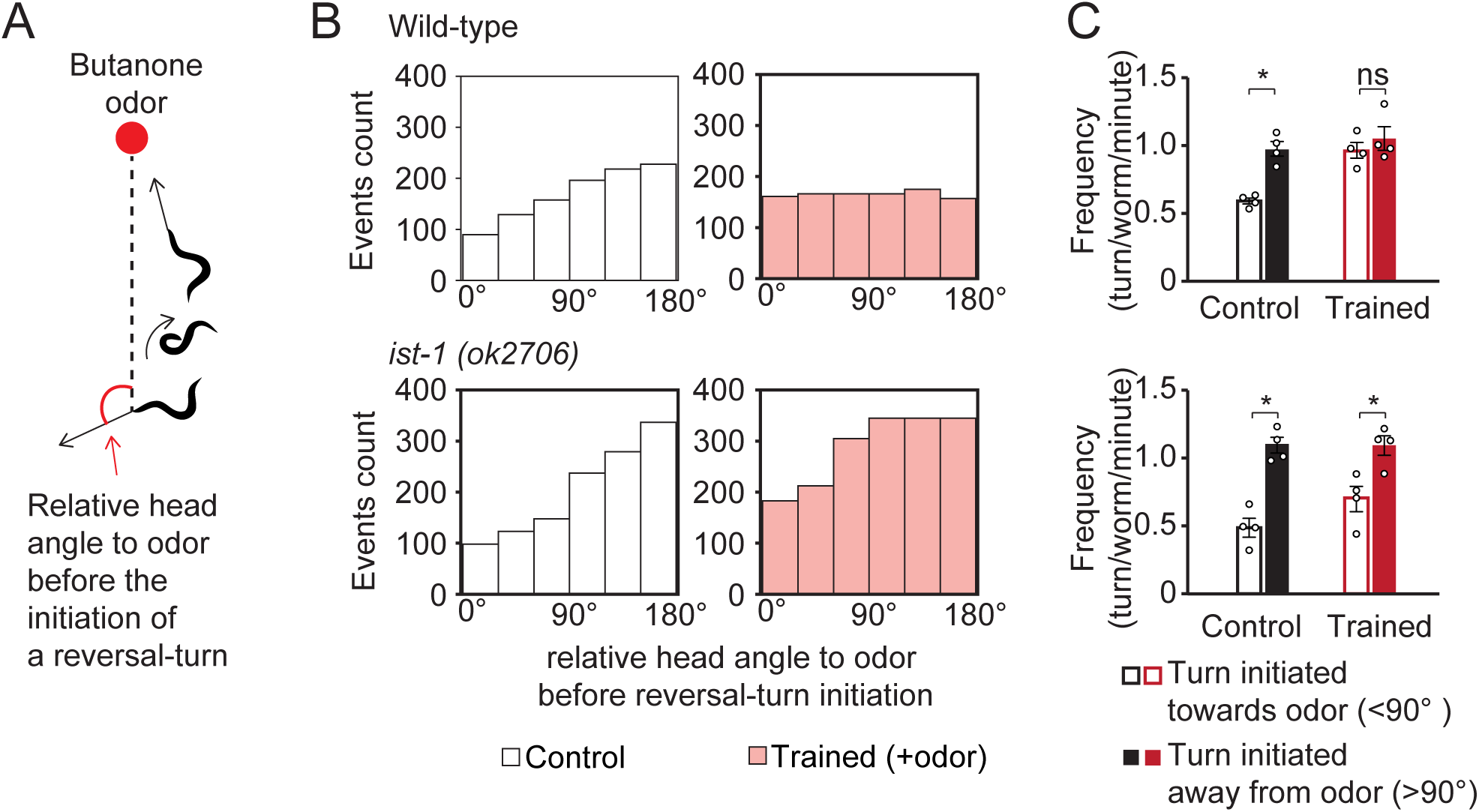
*ist-1* regulates biased random walk chemotaxis during aversive learning. A) *C. elegans* utilizes a biased-random walk strategy to navigate toward an odor source, in which turning increases as odor decreases. For panels (B) and (C), the angle between the animals’ heading direction and the direction of the odor source was determined shortly before each reversal-turn behavior during chemotaxis assays. B) Event count of reversal-turns for 50 wild-type or 50 *ist-1* mutant animals during one chemotaxis session. The angle between animals’ heading direction and odor source is binned by an increment of 30°. Animals were food-deprived (control, white bars), or food-deprived with butanone for 90 mins (trained, pink bars). C) Average frequency of reversal-turns for wild-type (black) or *ist-1* mutant (red) animals when they are traveling towards the odor (white bars with black and red borders), or traveling away from the odor (solid black and red bars). In (B) and (C), wild-type and *ist-1* control animals initiated more reversal-turn reorientations when traveling away from the odors, an act of course-correction for chemotaxis. Wild-type animals that were trained with butanone for 90 mins initiated similar frequencies of reversal-turns regardless of their heading direction, a finding that correlates with decreased chemotaxis after training. *ist-1* animals that were trained with butanone for 90 mins continued to initiate more reversal-turn reorientations when traveling away from the odors (*, values differ at p ≤ 0.05). Statistical tests are described in Table S2.

**Figure S2.**
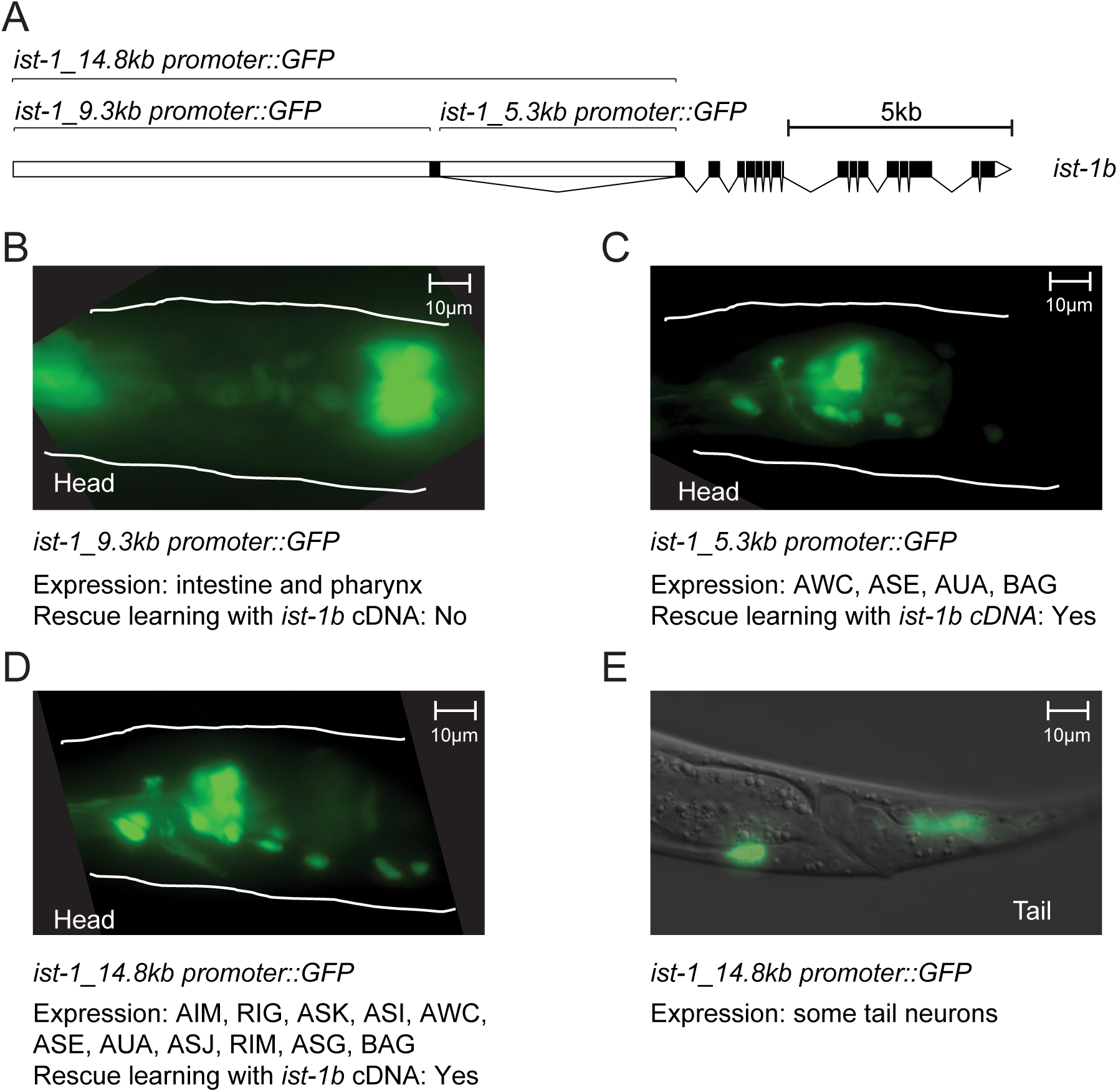
*ist-1* promoters and isoforms. A) Genomic structure of the *C. elegans* insulin receptor substrate-1 *(ist-1)* gene and flanking regions. White blocks indicate noncoding regions; black boxes indicate *ist-1* exons. B) The 9.3 kb genomic region between the first exon and the next gene upstream from *ist-1* drives GFP in the intestine and pharynx, and does not rescue *ist-1* aversive learning when driving an *ist-1b* cDNA. C) The 5.3 kb genomic region between the first exon and second exon of *ist-1* gene drives expression in the AWC, ASE, AUA and BAG neurons, and rescues *ist-1* aversive learning when driving an *ist-1b* cDNA. D-E) The 14.8 kb full genomic region drives expression in (D) AIM, RIG, ASK, ASI, AWC, ASE, AUA, ASJ, RIM, ASG and BAG neurons (Figure 2D) and (E) tail neurons, and rescues *ist-1* aversive learning when driving an *ist-1b* cDNA.

**Figure S3.**
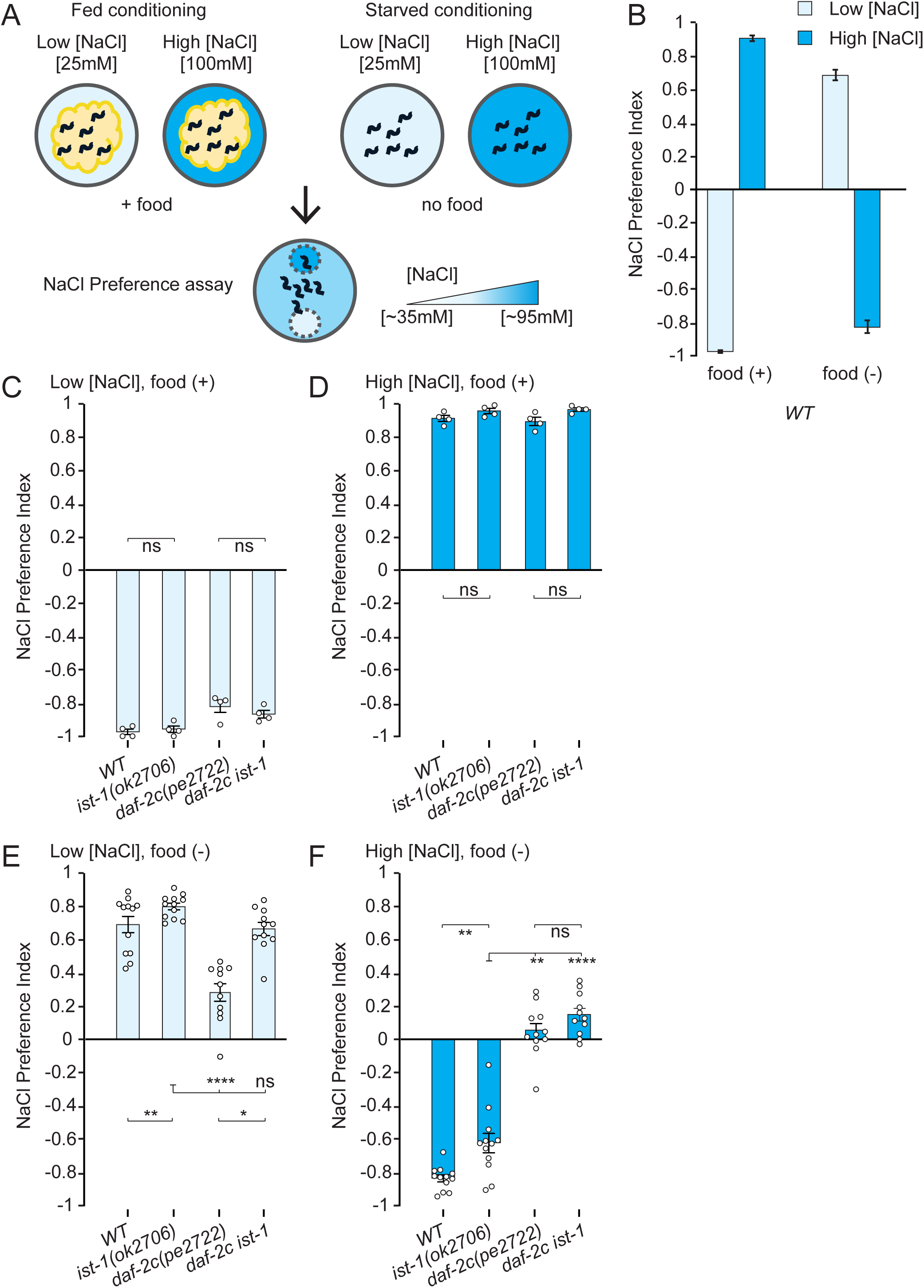
*ist-1* phenotypes and interactions with the insulin/IGF pathway in taste learning. A) Training and testing conditions for taste learning. Associative learning can be performed by pairing food or food deprivation with low (25 mM) or high (100 mM) salt concentrations. In the final salt preference assay, animals choose between low (35 mM) and high (95 mM) concentrations in a salt gradient. B) Salt preference after pairing food with low salt (light blue bars) or high salt (dark blue bars), or after pairing food deprivation with low salt (light blue bars) or high salt (dark blue bars). Animals prefer salt concentrations experienced with food. C-F) Salt preference of wild-type, *ist-1*, *daf-2c*, and *ist-1 daf-2c* double mutant animals after different training conditions. Animals were trained for 5 hours. Error bars indicate S.E.M. *, values differ at p ≤ 0.05. **, values differ at p ≤ 0.01. ****, values differ at p ≤ 0.0001. ns, values are not significantly different. Statistical tests are described in Table S2.

